# Toward a privacy-preserving predictive foundation model of single-cell transcriptomics with federated learning and tabular modeling

**DOI:** 10.1101/2025.01.06.631427

**Authors:** Jiayuan Ding, Jianhui Lin, Shiyu Jiang, Yixin Wang, Ziyang Miao, Zhaoyu Fang, Jiliang Tang, Min Li, Xiaojie Qiu

## Abstract

The ability to pre-train on vast amounts of data to build foundation models (FMs) has achieved remarkable success in numerous domains, including natural language processing, computer vision, and, more recently, single-cell genomics—epitomized by GeneFormer, scGPT, and scFoundation. However, as single-cell FMs begin to train on increasingly large corpora, significant privacy and ethical concerns arise. Moreover, unlike text data, single-cell data is unordered and exhibits a unique tabular structure that most existing single-cell FMs overlook. In this study, we propose Tabula, a privacy-preserving and tabular-structure aware FM designed with federated learning (FL) and tabular modeling. Tabula combines the advantages of FMs and FL, enabling collaborative model training across multiple clients without compromising data privacy. In contrast to earlier single-cell FMs—which treat single-cell data like natural language (viewing cells as “words” defined by genes)—Tabula introduces a novel pretraining strategy that explicitly models the tabular structure of single-cell data. Extensive experimental results show that Tabula outperforms state-of-the-art methods in various downstream tasks (including cell type annotation, gene imputation, gene perturbation, multi-batch integration, and multi-omics integration) while requiring only half the data for pretraining and preserving data privacy. Furthermore, Tabula accurately reveals pairwise and even combinatorial regulatory logic across diverse biological systems, including hematopoiesis, pancreatic endogenesis, neurogenesis, and cardiogenesis. Thus, Tabula provides a new foundation model that explicitly incorporates the tabular nature of single-cell data alongside FL, paving the way for creating a “virtual cell” for human health under critical privacy preservation.

## Introduction

One remarkable mystery in mutlicellular system is how the same genome can generate a vast diversity of cell types and states during development and diseases. Distinct gene expression patterns and complex pairwise or higher-order regulatory interactions underlie such diversities. Imagine each cell state as a point in the transcriptomic state space, where each dimension corresponds to a gene. The feasible space of cell states in any organism doesn’t occupy the entire state space but is generally considered to lie on a manifold, shaped by the governing regulatory networks.

Although genome-wide measurement of regulatory networks remains largely elusive, rapid advances in single-cell genomics and collaborative consortium efforts, such as CZI, the Human Cell Atlas (Rozenblatt-Rosen et al. 2017), and HuBMAP (HuBMAP Consortium 2019), have enabled us to map a significant portion of this single-cell manifold across human biology. Recent breakthroughs in natural language processing (NLP) (Devlin et al. 2019; Touvron et al. 2023; OpenAI et al. 2023), computer vision (Ranftl, Bochkovskiy, and Koltun, n.d.; He et al., n.d.), and, more recently, protein language models (Abramson et al. 2024; Lin et al. 2023; Rives et al. 2021) have shown that attention-based Transformers (Vaswani et al. 2017) can learn the fundamental structure of data through unsupervised self-learning on large datasets, rather than relying on explicit modeling of grammar, visual geometry, biophysical forces. The accumulation of the massive single cell data thus promises a new era of human biology, enabling the development of single cell foundation models to gain fundamental understanding of gene regulation—critical for virtually all cellular processes.

Over the past year, we have witnessed the emergence of single cell foundation models, notably Geneformer (Theodoris et al. 2023), scFoundation (Hao et al. 2024), CellPLM (Wen et al. 2023), scGPT (Cui et al. 2024), UCE (Rosen et al. 2023), and NicheFormer (Anna C. Schaar et al. 2024). However, current models fail to explicitly account for the tabular structure of the scRNA-seq data, often naively converting gene expression within a single cell to gene sequences to mimic word sequences used in NLP. While this adapts the NLP paradigm to single cell data, it overlooks critical features of the underlying data structure of scRNA-seq. Therefore, there is an urgent need to develop novel pretraining approaches to explicitly model the tabular structure of single cell data to enhance the foundation model pretraining. In addition, it has been shown previously that scRNA-seq data is susceptible to privacy breaches through linking attacks (Ferguson 2021; Walker et al. 2024), where information from one study can be linked to another to identify private data. This privacy concern becomes even more pronounced as we begin creating foundation models with tens of millions of single cells and datasets from thousands of individuals.

To address these critical gaps, we developed Tabula, a privacy-preserving federated foundation model tailored for scRNA-seq data through tabular learning. Tabula delivers two major innovations. First, it introduces federated learning (FL) (Kairouz et al. 2021) into the foundation model training, allowing for collaborative model training across multiple clients without compromising the privacy of the individual data. The distributed computing framework also significantly reduces local training cost. Furthermore, while Tabula’s FL framework relies on a shared tabular transformer to harbor common knowledge across clients, it also possesses client-specific embedders and project heads to capture unique characteristics of each client, e.g. tissue. This special design improves zero-shot performance on downstream tasks specific for each client. Secondly, unlike earlier single cell foundation models (Cui et al. 2024; Theodoris et al. 2023; Hao et al. 2024; Yang et al. 2022), which treated single cell data analogous to natural language data (e.g., viewing cells as sentences and genes as words), Tabula redefines this paradigm by treating single cell data as tabular data and modeling it accordingly with tabular learning. Extensive benchmarking demonstrates that Tabula consistently outperform state-of-the-art foundation models on gene-wise and cell-wise downstream tasks, including cell type annotation, batch correction, gene imputation, single- and double-perturbation response prediction and reverse perturbation predictions. Importantly, we also show that Tabula can reveal pairwise and even combinatorial gene regulations across diverse biological systems, including hematopoiesis, pancreatic endogenesis, neurogenesis and cardiogenesis where extensive biochemical studies have validated core regulatory networks. These results represent a significant step toward building a mechanistic and predictive single cell foundation model. Tabula, part of our AI agent ecosystem Chiron, is implemented with Pytorch Lightning and freely accessible at https://github.com/aristoteleo/tabula.

A decade ago, with the advent of next-generation sequencing of the human pathogen Mycoplasma genitalium, Karr et al. (Karr et al. 2012) reported the first whole-cell model that synthesizes diverse mathematical approaches to predict a broad spectrum of biological processes. Given the recent advancements in single-cell genomics, along with ever-expanding cell atlas profiling efforts, the creation of virtual cells—a long-sought goal in computational biology—seems increasingly attainable. However, instead of relying on explicit biophysical models as demonstrated by Karr et al., advances in generative AI will drive this new phase of virtual cell development. Generative AI has shown incredible promise in understanding multi-scale and complex patterns across domains such as natural language processing (NLP) and computer vision, similar to biology where modeling biological systems with thousands of variables, especially in mammalian systems, is unfathomable. Since an ultimate single cell foundation model could theoretically synthesize all known knowledge of human biology, we envision a future where the virtual cell based on the single cell foundation model can be used to simulate every aspect of cell development, diseases, reshaping the landscape of human medicine by predicting the optimal drug, gene, and cell therapies that revert the disease states back to the normal.

## RESULTS

### Tabula redefines single cell foundation models through tabular and federated learning

Pioneer single cell foundation models such as Geneformer (Theodoris et al. 2023), scGPT (Cui et al. 2024) show promises of LLM in advancing single cell biology. However, these frameworks and recent new developments, e.g. UCE (Rosen et al. 2023), CellPLM (Wen et al. 2023), NicheFormer (Anna Christina Schaar et al. 2024) etc., rely on centralized learning that trains a unified model directly on aggregated data, thus raising concerns about data privacy and ethics (Ferguson 2021; Walker et al. 2024). Furthermore, although it is obvious that single cell data doesn’t have an intrinsic order, most of these approaches nevertheless directly borrow conventional large language model techniques originally designed for sequential text data to single cell data after transforming the unordered genes across single cells to gene sequences with some rather naive ordering strategies **(Fig. 1a)**. For example, Geneformer generates a gene pseudo-order by ranking the normalized gene expression within each cell, followed by masked language modeling (Devlin et al. 2019). Similarly, scGPT first bins genes and then uses attention maps from the learned transformer to create a sequence order by prioritizing genes with high attention scores for next-token prediction using autoregressive modeling (Hsiao and Hung 2022). Likewise, tGPT (“Generative Pretraining from Large-Scale Transcriptomes for Single-Cell Deciphering” 2023) creates a pseudo-order of genes by sorting the normalized gene expression within each cell and employs autoregressive modeling to predict the subsequent gene token.

**Figure 1:**
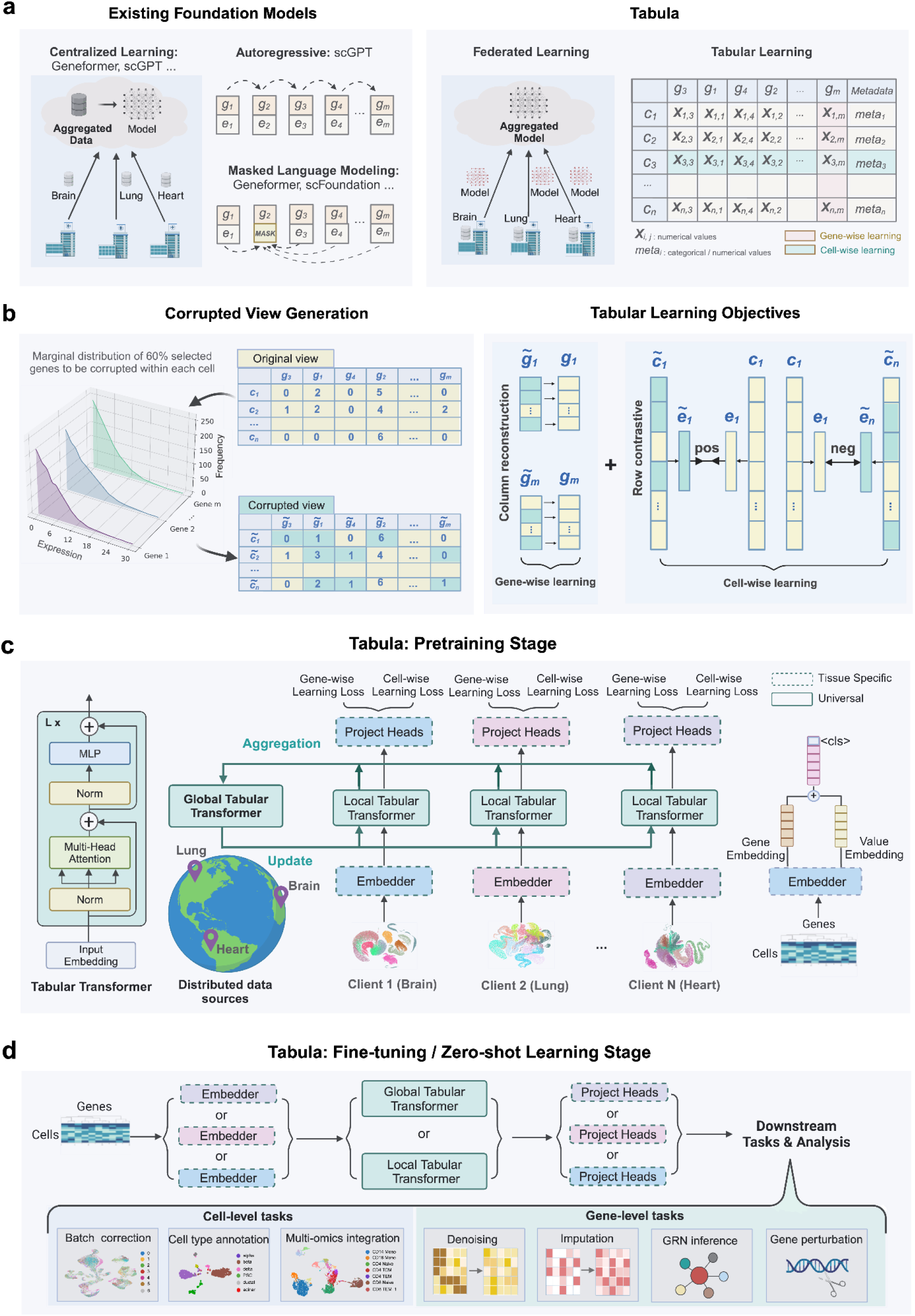
The schematic of Tabula, a novel single cell foundation model that integrates federated learning and tabular learning. **a. Key conceptual innovations of Tabula over existing single cell foundation models**. While existing foundation models, e.g., Geneformer (Theodoris et al. 2023) and scGPT (Cui et al. 2024) rely on centralized models that train on the compiled corpus of shared data on a central server, Tabula employs federated learning (Kairouz et al. 2021), independently on decentralized clients avoiding sharing raw data. Furthermore, although gene expression from single cells don’t have a natural order, existing models treat gene expression within each cell from the scRNA-seq data as ordered gene sequences and use autoregressive or masked language modeling for unsupervised learning, following the conventional paradigm in large language models for text data. Instead, Tabula explicitly considers the tabular structure of scRNA-seq data and uses tabular learning to build the foundation model. In Tabula, each cell is represented by a permutation-invariant row of genes (each gene corresponds to a column) containing numerical expression values or metadata categories, respecting the unordered structure of gene features. **b. Tabula relies on self-supervised tabular modeling of corrupted expression data along both gene and cell dimensions**. Tabula’s self-supervised tabular learning starts with creating a corrupted view of the scRNA-seq data by resampling a subset of genes and replacing the expression values based on their individual gene column marginal distribution. For instance, *g*_3,_ *g*_1,_ *g*_2_ are randomly selected and replaced with the samples values 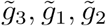 from their corresponding marginal distributions in. *c*_1_ Similarly, a different set of 60% of genes including *g*_1,_ *g*_4,_ *g*_*m*_ are corrupted in *c*_*n*_. Tabula then applies two objectives: gene-wise (column) learning, which reconstructs the original gene view from its corrupted gene view, and cell-wise (row) learning, which uses contrastive learning on the latent space to distinguish between positive and negative cell pairs. This modeling strategy captures the structure of single cell data along both the gene and cell axis. **c. The model architecture and pretraining scheme of Tabula**. Tabula uses a transformer (Vaswani et al. 2017) for both local or global modeling. During pretraining, Tabula’s local model at each client is updated independently. Then the updated local model weights are sent to a central server without sharing the raw data, thus preserving data privacy. In this study, we design each client specific for a tissue type (e.g., lung, brain, heart) that comprises a tissue-specific embedder and project heads that capture the tissue’s unique characteristics. Local training alternates with global model updates in a continuous privacy-preserving cycle across clients. **c. Downstream fine-tuning and zero-shot learning by Tabula**. After pretraining, Tabula utilizes its tissue-specific embedders and transformers (either local or global) for downstream tasks at the gene and cell levels. Cell-level tasks include batch correction, cell annotation, and multi-omics integration, while gene-level tasks cover denoising, imputation, gene regulatory network inference, and gene perturbation prediction.

To explicitly model the tabular structure while preserving data privacy, we propose Tabula, a novel single cell foundation model that integrates tabular and federated learning **(Fig. 1a)**. First, we leverage federated learning (Kairouz et al. 2021) to allow Tabula to have different clients and to train a client-specific model locally on private data without sharing raw data across institutions, thus eliminating privacy concerns for potential single cell datasets generated from patient or proprietary samples. Second, instead of treating gene expression data as ordered sequences, we explicitly model its intrinsic tabular structure using self-supervised tabular learning (Gorishniy et al. 2021). Specifically, we treat each cell of the gene expression matrix as a permutation-invariant row of genes, where each column corresponds to a specific gene (e.g., *g*_1_, *g*_2_, … *g*_*m*_), and both cell and genes involves numerical expression value, e.g., *X*_*i,j*_ representing the measured expression level of gene *j* in cell *i*. The column feature can also include metadata, such as tissue type or cell type, providing additional biological context. Tabula leverages a large-scale pretraining dataset comprising 15 million single cells across diverse tissues (**Supplementary Fig. 1a and b**), following a scaling law that demonstrates significant performance improvements as the pretraining dataset size increases (**Supplementary Fig. 1c and d**). Additionally, Tabula can generate tissue-specific embedders that encode distinctive features of individual tissues, allowing for better representation of tissue-specific biological characteristics (**Supplementary Fig. 2**). Unlike sequence-based models that impose an artificial order on genes, Tabula’s tabular design treats each gene as an independent attribute, respecting their unordered nature. By incorporating both cell-wise and gene-wise learning (**Methods**), Tabula effectively models both gene and cell relationships of the tabular single cell data. These innovative designs in Tabula necessitate new training loss for tabular data, as well as model architecture and pretraining scheme.

To handle the intrinsic tabular structure of scRNA-seq dataset, Tabula introduces an innovative self-supervised tabular modeling framework that consists of both gene-wise reconstruction learning and cell-wise contrastive learning (**Fig. 1b**). Both training components start with generating a corrupted representation of the scRNA-seq data. For gene-wise reconstruction, 60% of genes in each cell are firstly randomly selected, expression of these selected genes are replaced with values sampled from each gene’s marginal distributions across cells (**Methods**). Then gene-wise reconstruction tries to restore original expression values (columns of the *cell by gene* table) from their corrupted views. In contrast, the cell-wise contrastive learning tries to align or separate positive or negative cell pairs in latent spaces, respectively. Specifically, the gene expression data of each cell, including original (*c*_1_) and corrupted views 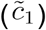, is firstly embedded into latent dimension to obtain *e*_1_ and 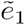. Then contrast learning is used to ensure positive cell pairs (e.g., *e*_1_ and 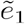), representing the same cell, colocalize while negative pairs (e.g., *e*_1_ and 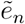, two different cells), representing distinct cells, overall separate in the latent space. Therefore, while gene-wise reconstruction ensures robust gene-level representations, the cell-wise contrastive learning ensures both intra-cellular consistency but also inter-cell variability, thus properly modeling the tabular structure of single cell cell-by-gene matrix along both the gene and cell axis.

Additionally, to address the data privacy concern, Tabula integrates federated learning that relies on the distributed training across clients without sharing raw data (**Fig. 1c**). Tabula’s federated learning framework builds upon a unified transformer with the architecture shared between both the client model and global model but trained separately. Tabula’s pretraining leverages large-scale, privacy protected datasets, each tied to a specific client, such as a hospital, institute or tissue type (e.g., lung, brain, heart), as demonstrated here. Raw data remains localized at client sites, while only local tabular transformer weights are shared with a central server to update the global tabular transformer model. Each client, containing a client-specific embedder and project heads (**Fig. 1c**), transforms cell-by-gene tables into embeddings, which are processed by the tabular transformer to compute gene-wise and cell-wise learning losses. The training process involves iterative updates of the local and global tabular transformers in alternating cycles. At the client level, local transformers learn client-specific, e.g. tissue-specific, patterns which are then uploaded to the center server. At the center level, the global transformer aggregates these uploaded weights to update the global model and then broadcasts the aggregated weights back to local clients to build a more generalized model across clients. This continuous communication between the center server and the client ensures exchange of knowledge across diverse data sources without data sharing and thus preserving data confidentiality. By training on distributed clients using tabular learning loss, Tabula establishes a novel foundation model for single cell data that is privacy-preserving, capable of capturing tissue-specific and cross-tissue biological variations while respecting the intrinsic tabular structure of the scRNA-seq data.

Built upon these innovations in Tabula, we apply Tabula to a series of cell-level or gene-level fine-tuning and zero-shot learning applications (**Fig. 1d**). While cell-level tasks include batch correction, cell type annotation, and multi-omics integration, the gene-level tasks include imputation, GRN inference, and gene perturbation analysis. More importantly, as a testimony to prove that a single cell foundation model can achieve mechanistic and predictive insights, we firstly show that Tabula accurately predicts pairwise and high-order combinatorial regulation across a range of distinct biological systems. We further demonstrate that Tabula is able to accurately predict transcriptional responses to diverse perturbations and how genetic perturbations divert cell lineages.

Taken together, Tabula is a novel single cell foundation model that relies on tabular learning and federated learning to better model the intrinsic structure of the scRNA-seq data while ensuring privacy preservation, promising in gaining mechanistic and predictive insights, as described in the following.

### Tabular learning, a novel single cell pretraining strategy introduced in Tabula, outperforms masked language modeling (MLM) and autoregressive modeling (AR) in both federated and centralized learning settings

While pioneering single cell foundation models simply order gene expression within each cell to obtain gene sequences and then proceed with natural language approaches for centralized pretraining, Tabula explicitly models the matrix structure of scRNA-seq data with tabular learning for federated pretraining to preserve data privacy. To demonstrate the advantages of tabular learning and its adaptability to federated learning, we firstly implemented a well-controlled and systematic benchmark on two common cell-level or gene-level downstream tasks, naming cell-type annotation and genetic perturbation prediction, by enumerating all possible four combinations of pretraining framework (federated vs. centralized learning) and single cell modeling strategies (tabular learning vs. MLM) (**Fig. 2a**). Specifically, we performed benchmarks on cell-type annotation and gene perturbation prediction with all four training designs on a small-scale pretraining dataset comprising 1 million (M) human cells in total, with 250,000 cells sampled from each of four distinct tissues, pancreas, blood, brain, and lung, sourced from the CELLxGENE census (**Methods**). While the centralized training uses the aggregated 1M single cell dataset, the federated learning employs four clients, each initialized to be trained on the data subset from a particular tissue, with epoch-wise aggregation of Tabula transformer weights across clients. Both training frameworks were used in conjunction with either tabular learning (**Methods**) or masked language modeling (MLM) with a 15% masking ratio specific to MLM, resulting in four combinatory pretraining designs. To further demonstrate the advantages of tabular learning and federated learning, we next benchmarked the performance of the fully tabula model against state-of-the-art foundation models such as scGPT (Cui et al. 2024), scFoundation (Hao et al. 2024), and Geneformer (Theodoris et al. 2023) (**Fig. 2b**).

**Figure 2:**
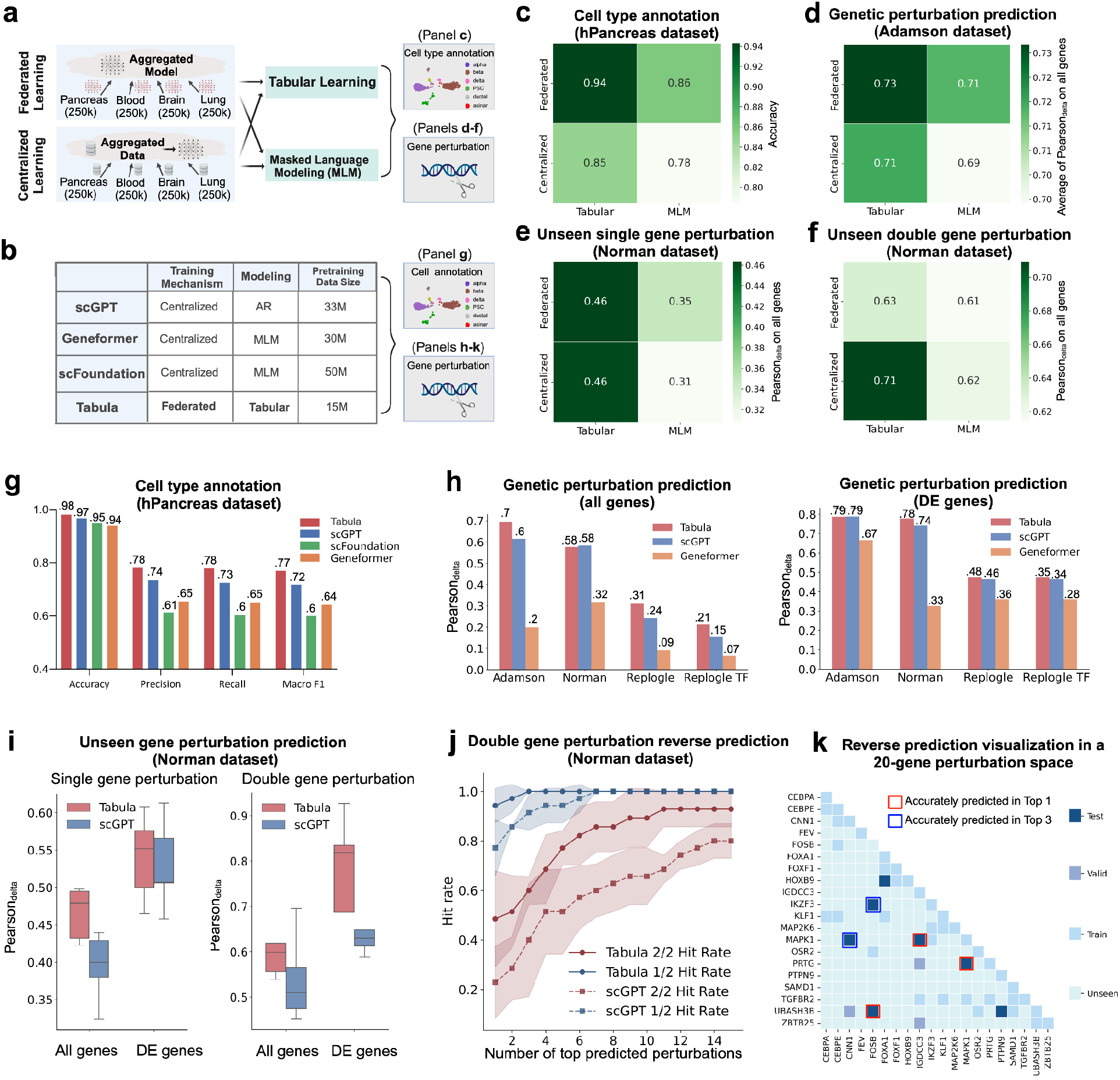
Pretraining with Tabular learning consistently outperforms that with masked language modeling (MLM) or autoregressive modeling (AR), while being generally applicable to both the federated and centralized learning settings. **a. Benchmark design for evaluating the performance of cell-type annotation and gene perturbation prediction with all combinations of training frameworks (federated vs. centralized learning) and single cell modeling strategies (Tabular vs. MLM learning)**. **b. Training mechanisms, modeling approaches and pretraining data size for scGPT, Geneformer, scFoundation and Tabula**. While Tabula is the only framework that uses federated learning, all other frameworks, including scGPT, Geneformer, and scFoundation rely on centralized learning. Furthermore, regarding pretraining strategies, scGPT employs autoregressive modeling, Geneformer and scFoundation are based on MLM, and Tabula adopts tabular learning. The pretraining data sizes also vary, with Tabula using the smallest dataset (15M cells) and scFoundation the largest (50M cells). Both cell type annotation and genetic perturbation prediction are used for downstream tasks. c. Heatmap illustrates cell type annotation performance (accuracy) on the hPancreas dataset (J. Chen et al. 2023) across four pretraining combinations: federated vs. centralized learning and tabular learning vs. masked language modeling (MLM). d. Same as in panel **c** but for the average Pearson correlation coefficient (Pearson_delta_) between predicted and observed gene expression changes across all genes on the single perturbation Adamson dataset, with higher values reflecting better alignment between predicted and observed gene expression changes (Adamson et al. 2016). e. Same in panel **c** but for the Pearson correlation coefficient for unseen single-gene perturbations on the Norman dataset (Norman et al. 2019). f. Same as in panel **e**, but for unseen double-gene perturbation on the Norman dataset (Norman et al. 2019). g. Tabula significantly surpasses scGPT, Geneformer and scFoundation in cell type annotation on the hPancreas dataset, achieving highest performance across all four metrics: Accuracy, Precision, Recall, and Macro-F1. **h. Tabula outperforms scGPT and Geneformer in predicting gene expression changes following genetic perturbations**. Barplots of Pearson correlation coefficients (Pearson_delta_) between predicted and observed gene expression changes for each model are shown across single perturbation datasets (Adamson (Adamson et al. 2016), Replogle (Replogle et al. 2022) and Replogle TF (Replogle et al. 2022) datasets) and double perturbation Norman dataset (Norman et al. 2019). The left subpanel shows correlation coefficients for all genes, while the right subpanel focuses only on differentially expressed (DE) genes. Higher Pearson correlations reflect greater accuracy in capturing true perturbation effects. **i. Tabula outperforms scGPT in predicting expression changes of unseen single and double-gene perturbation in the Norman dataset** (Norman et al. 2019). Random dataset splits across five seeds are used for both Tabula and scGPT model predictions. The left subpanel illustrates Pearson correlation coefficients for unseen single-gene perturbations, while the right subpanel depicts correlations for unseen double-gene perturbations (both genes are unseen or unused during the fine-tuning stage). Box plots present the distribution of Pearson correlations for all genes and the top 20 differentially expressed (DE) genes, with higher values indicating closer alignment between predicted and observed gene expression changes. Tabula demonstrates overall superior performance over scGPT, particularly in double perturbation settings. Here the central line of the boxplot indicates the median and the box edges representing the first (Q1) and third (Q3) quartiles, where whiskers extend to the most extreme data points within 1.5 times the IQR, while individual points beyond the whiskers denote potential outliers. **j. Hit rate analysis for reverse perturbation predictions of two-gene combinations**. The lineplot shows hit rates for the top *K* predicted perturbation combinations evaluating forTabula and scGPT on five random seeds. Lines represent the average hit rate of each model in predicting one of two genes correctly (1/2 Hit Rate) or both genes correctly (2/2 Hit Rate). The transparent shade indicates the 95% confidence interval. As *K* increases, both models show improved hit rates, with Tabula generally outperforming scGPT in both 1/2 and 2/2 hit rates, indicating greater accuracy in reverse identifying correct gene perturbation combinations. **k. Heatmap of reverse dual-perturbation prediction across the all possible 20-gene pairwise perturbation space**. The grid represents the perturbation combination from genes on the row or column, while filled color for each grid represents different computational modeling data types, including train, validation (valid), test, and unseen data. Predicted perturbation combinations are highlighted by square boxes, with red squares indicating the combinations accurately predicted as the first perturbation combinations and blue squares for those accurately predicted among the top 3 perturbation combinations.

Our evaluation reveals that Tabula learning consistently outperforms MLM in both federated and centralized learning settings in annotating cell type and predicting genetic perturbation response. For the cell type annotation benchmark, the hPancreas dataset (J. Chen et al. 2023) was used (**Downstream Task Datasets**). Notably, federated learning outperformed centralized learning, and tabular learning consistently demonstrated superior performance compared to MLM (**Fig. 2c**). For genetic perturbation prediction, we first assessed performance on the Adamson (Adamson et al. 2016) single-gene perturbation dataset (**Downstream Task Datasets**) by averaging the Pearson correlation coefficients between predicted and observed gene expression changes (*Pearson*_*delta*_) over all genes (**Fig. 2d**). Subsequently, we evaluated the Norman double-gene perturbation dataset (Norman et al. 2019), focusing on the *Pearson*_*delta*_ for all genes, including cases of unseen single- and double-gene perturbations (**Fig. 2e and f**). Across these evaluations, Tabular learning consistently outperformed MLM, and federated learning maintained performance comparable to centralized learning. These results underscore the robustness of tabular learning in capturing informative cell-level representations for tasks like cell type annotation, while also excelling in gene-level tasks, such as genetic perturbation prediction, by effectively modeling gene-wise relationships.

Have demonstrated the improvement of tabular learning over MLM from the small-scale models of the benchmark framework we implemented, we now demonstrate how our final pretrained Tabula model performs against state-of-the-art foundation models such as scGPT (Cui et al. 2024), scFoundation (Hao et al. 2024), and Geneformer (Theodoris et al. 2023). The Tabula model was pretrained on 15 million cells from the CELLxGENE census using a federated learning framework optimized through tabular modeling (**Methods**). In contrast, the other three models employed centralized pretraining with a natural language modeling paradigm and utilized training datasets at least twice the size of that of Tabula. For cell type annotation, Tabula outperformed scGPT across all four metrics—accuracy, precision, recall, and F1—while scFoundation and Geneformer demonstrated even poorer performance, despite all three models being pretrained on datasets at least twice the size of Tabula’s (**Fig. 2g**) (**Supplementary Fig. 3**). Next, we conducted benchmark experiments on genetic perturbation prediction across four datasets: Adamson (Adamson et al. 2016) and Replogle (Replogle et al. 2022) for single-gene perturbation, Norman (Norman et al. 2019) for double-gene perturbation, and Replogle TF (Replogle et al. 2022) for transcription factor-specific perturbation (**Fig. 2h**). Using *Pearson*_*delta*_, the Pearson correlation between the average predicted and observed changes across cells after gene perturbation as a benchmark metric, Tabula outperformed scGPT on the Adamson dataset (“all-gene” setting or when all genes are used for the benchmark) and on the Norman dataset (“DE-gene” setting when only top 20 differential expressed genes based on observed data are used for the benchmark), while achieving comparable performance to scGPT on the Adamson dataset in the DE-gene setting and the Norman dataset in the all-gene setting. Tabula demonstrates superior results on the Replogle dataset on both settings. By contrast, Geneformer consistently underperformed relative to the other models.

We further investigated Tabula’s capabilities on predicting unseen genetic perturbation responses and recovering causal gene pairs from gene expression signatures after perturbations. To rigorously assess the performance of fine-tuned models on *in silico* prediction of genetic responses to unseen perturbations, we conducted five independent experiments with random seeds, using unseen single- and double-gene perturbation cases during fine-tuning from the Norman dataset (Norman et al. 2019) (**Fig. 2i**). While scGPT performs comparably to Tabula on the top 20 differentially expressed genes in single-gene cases, Tabula significantly surpasses it in all other scenarios, underscoring its overall improvement in the perturbation prediction compared to scGPT. To evaluate Tabula’s effectiveness in recovering causal gene pairs from gene expression signatures after perturbations, we followed the experimental setup of scGPT, focusing on a subset of the Norman dataset (Norman et al. 2019), which comprises perturbations involving 20 genes and 210 possible one-gene or two-gene combinations. To ensure fair and robust evaluation, we performed five iterations of random data splitting into training, validation, and test sets, with proportions of 75%, 15%, and 10%, respectively, following the setting of GEARS (Roohani, Huang, and Leskovec 2023). The model’s predictions were assessed based on their rank within the top predictions to measure its ability to recover the correct genetic perturbations associated with given cell states subject to specific genetic perturbations. Tabula demonstrated a remarkable 2/2 hit rate of 48.57% and a 1/2 hit rate of 94.29% when evaluating the top 1 predicted perturbation combinations, substantially outperforming scGPT (**Fig. 2j**). Additionally, we further demonstrated that Tabula is able to accurately pinpoint genetic perturbations for the majority of test cases within the 20-gene combinatorial perturbation space (**Fig. 2k**). For instance, perturbations such as FOSB+UBASH3B and CNN1+MAPK1 were successfully ranked as the top 1 and top 3 predictions, respectively, highlighting Tabula’s capability in identifying critical gene combinations driving cell state transitions. These results support Tabula’s superior performance in reverse perturbation prediction, making it a valuable tool for accelerating the discovery of genetic drivers for development and disease as well as optimizing experimental perturbation designs. Extensive evaluation across perturbation and reverse-perturbation prediction tasks, including single- and double-perturbation settings, diverse experimental contexts, and varying gene expression levels (**Supplementary Fig. 4**), further demonstrates Tabula’s superior accuracy in unveiling complex inter-genetic relationships. Additionally, we benchmarked Tabula against state-of-the-art foundation models on extra gene-level or cell-level downstream tasks: imputation (**Supplementary Fig. 5**), multi-omics and multi-batch integration (**Supplementary Fig. 6**). Tabula outperformed all of them across these tasks.

In conclusion, our findings consistently demonstrate that Tabula achieves favorable performance across both gene-wise and cell-wise downstream tasks, as evidenced by experiments comparing federated versus centralized learning and Tabular versus MLM learning as well as predictions of single- and double-perturbation response and reverse predictions. These results underscore the effectiveness of Tabula’s unique tabular modeling approach that explicitly accounts for the tabular structure of scRNA-seq, enabling favorable performances over state-of-the-art approaches, despite the use of limited pretraining data.

### Tabula accurately recovers pairwise and high-order combinatorial regulation across diverse biological systems with zero-shot prediction

Although the single cell foundation model has shown to be promising in many different tasks, a true testimony of a valuable single cell foundation model should prove its accuracy, consistency and generalizability when applied under zero-shot setting, e.g., completely new situations without prior specific training, and should confirm its ability in deriving mechanistic biological insights from a very diverse set of biological systems. To demonstrate this, we take on a novel and challenging task by validating Tabula’s performance in recovering pairwise and higher-order combinatorial gene regulation across four distinct developmental systems, ranging from hematopoiesis (Qiu et al. 2022; Krumsiek et al. 2011), cardiogenesis (de Soysa et al. 2019; Nakanishi et al. 2016), neurogenesis (La Manno et al. 2021; Qiu et al. 2012), and pancreatic endogenesis (E17) (Wang et al. 2023; Wang et al. 2020), that have accumulated very well validated regulatory networks through decades of biochemical research.

To enable Tabula to predict pairwise and combinatorial regulations, we proposed a novel strategy based on zero-shot prediction. For the pairwise regulation prediction, we iteratively in silico perturb the expression of a regulator gene, e.g., *g*_1_, with our tabula model, while tracking the resulting impact on its target (*g*_2_) over multiple iterations (**Fig 3. a**). Then the dynamic trajectories of both the regulator and target will be characterized, which is further used to determine activation, repression, or neutrality regulation mode based on expression distribution shifts (**Fig. 3a-iv**) (**Fig. 3a-v**) of the target before and after the iterative perturbation, as well as the proportion pattern of different regulation modes across cells. For example, in the illustrated diagram, *g*_2_ expression decreases after in silico activating *g*_1_ expression, thus indicating a repression from *g*_1_ to *g*_2_. A combinatorial regulation involving multiple regulators to the same target can be often simplified as a boolean table. Therefore, for the combinatorial regulation predictions, Tabula first enumerates all possible boolean logic gates of the combinatorial regulation and then follows the procedure from pairwise regulation prediction to perform iterative perturbation prediction based on the logic gate. Specifically, if the value of a regulator is set to be zero or one on the logic gate, the regulator will decrease or increase the gene rank by one. For example, we show Tabula is able to accurately predict the combinatorial regulation from GATA1 and FLI-1 to EKLF as it only increases EKLF’s gene expression after we increase GATA-1’s expression order by 1 while decreasing FLI-1’s order by 1 iteratively but not others. This simple but powerful strategy provides a general approach for Tabula to predict complex gene regulatory scenarios.

**Figure 3:**
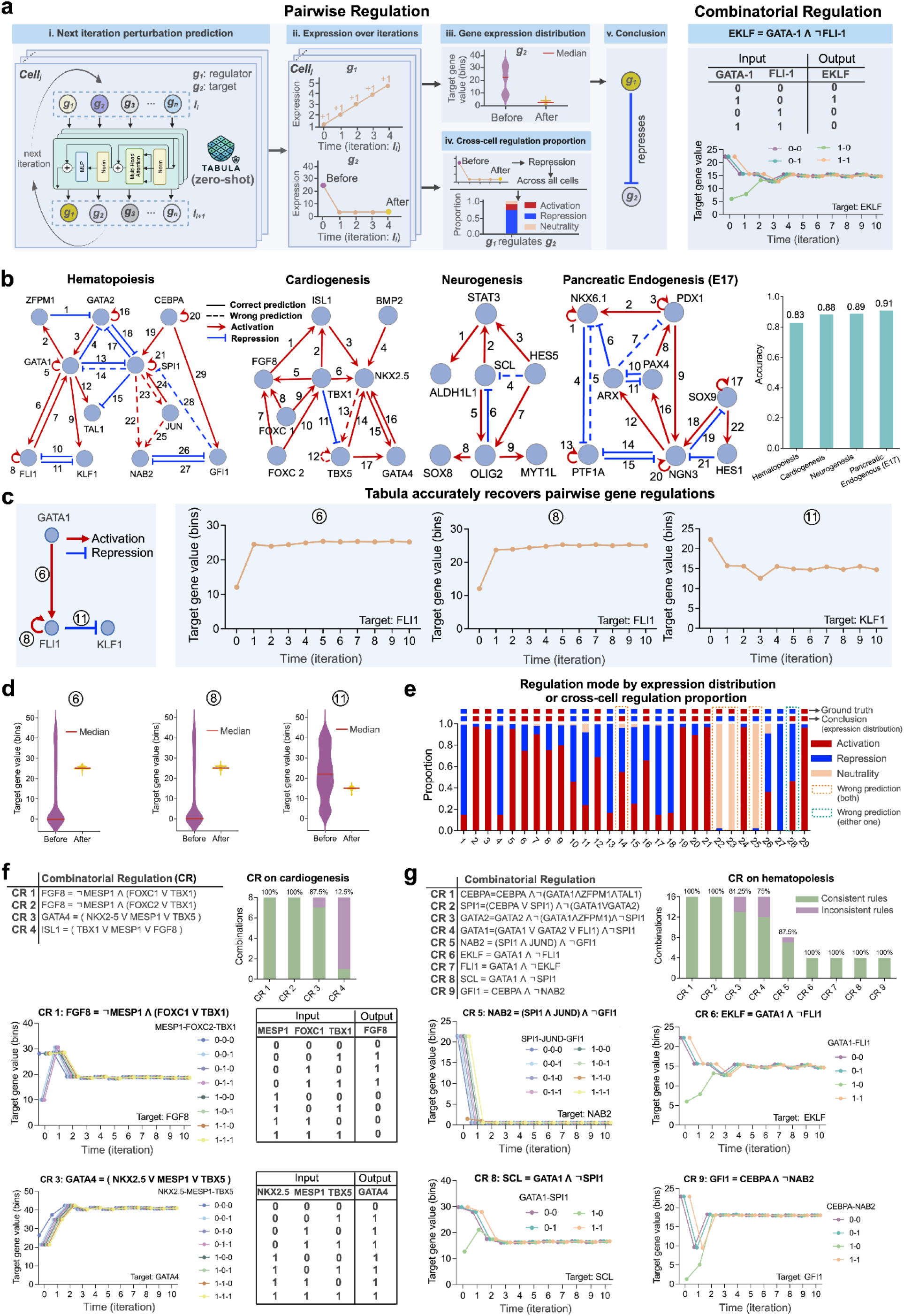
Tabula accurately predicts pairwise and combinatorial regulation across a variety of biological systems. **a. The schematic of utilizing Tabula zero-shot prediction to characterize both pairwise and combinatorial regulations**. The left subpanel outlines Tabula’s strategy for predicting pairwise regulation across cells between a regulator *g*_1_ and its *g*_2_ target through an iterative perturbation zero-shot prediction procedure. (i) This iterative update procedure involves consistently increasing or decreasing the rank of gene expression of the regulator by one across cells, followed by zero-shot predicting the expression of all genes after passing through the Tabula model. Then the updated gene expression is used as the input for the next iteration. (ii) The predicted trajectories of the expression dynamics across both the regulator and target under the above *in silico* perturbations can then be obtained and subsequently used to reveal *g*_1_’s effect on *g*_2_ (iii) Diagram of target gene expression distribution. Expression distribution (median values indicated by red lines) of the target before and after the iterative perturbation. By default, this approach is used to conclude the type of gene regulation, see (v). (iv) Diagram of cross-cell regulation proportion. Cross-cell regulation proportions depict the relative fraction of activatory, repressive, or neutral regulation modes from the regulator to the target across all cells. (v) Based on the median gene expression before and after iterative perturbation, we can conclude the gene regulation mode. In this case, we find the expression of *g*_2_ decreases after the iterative perturbation and we thus conclude a repression regulation from *g*_1_ to *g*_2_ Both the median expression based regulatory mode and cross-cell regulation proportion can be used for determining the gene regulation mode in this study. The right subpanel illustrates how Tabula determines combinatorial regulation using the example of the combinatorial regulation from GATA1 and FLI-1 to EKLF. We first enumerate all possible boolean logic gates from GATA1 and FLI1 to EKLF and follow the procedure from the left panel to perform iterative perturbation prediction where the perturbation is executed on both the GATA1 and FLI1. Note that if the value is set to be zero (one) on the logic gate, the regulator will decrease (increase) the gene rank by one. Similarly, based on the target gene expression dynamics across iterations under different perturbation input, we conclude it matches up with the combinatorial logic from GATA1 and FLI1 to EKLF. **b. Four biological systems with extensively validated regulatory networks that Tabula applied to and its overall performance on pairwise regulation prediction**. The core regulatory networks for hematopoiesis (Fig. 4b-e) (Krumsiek et al. 2011), cardiogenesis (Fig. 4b) (Nakanishi et al. 2016), neurogenesis (Fig. 4b) (Qiu, Ding, and Shi 2012), and pancreatic endogenesis (E17) (Fig. 4b) (J. Wang et al. 2020) is shown on the left. In the network diagram, an activation regulation is denoted by red arrows, repression regulation by blue arrows, with solid arrows indicating correct predictions and dashed arrows incorrect ones. The bar plot on the right quantifies the global performance of the Tabula model across all systems. c. Tabula recovers pairwise gene regulations within a three-gene example network motif in the hematopoiesis system (Krumsiek et al. 2011). The left diagram illustrates regulatory relationships between GATA1, FLI1, and KLF1. The line plots on the right display predicted expression levels of target genes (FLI1 and KLF1) over iterations after increasing the regulator, matching up with expected expression dynamics. **d. Distribution of target gene expression pre- and post-perturbation under Tabula prediction in a three-gene example network motif from the hematopoiesis system** (Krumsiek et al. 2011). Violin plots illustrate expression levels of target genes for interactions 6, 8, and 11 before and after perturbation. **e. Comparison of regulation mode identified by expression distribution shift and cross-cell regulation proportions**. Bar plot shows activation (red), repression (blue), and neutrality (orange) proportions for each interaction of the hematopoiesis system (Krumsiek et al. 2011), based on cross-cell regulation proportions (see panel **a-iv**). The first row of the regulation mode shown on the top of each bar represents ground truth mode while the second row of the regulation mode indicates the prediction from the expression distribution. Highlighted boxes with dashed lines denote inconsistent prediction: orange dashed box for incorrect predictions from both expression distribution shift based and cross-cell regulation proportion based regulation modes, while blue dashed box for incorrect prediction in only one such prediction. **f. Combinatorial regulation (CR) predictions for the cardiogenesis system** (Herrmann et al. 2012). The table (**top left**) lists four CR rules, where multiple regulators work in concert to regulate the expression of FGF8, GATA4, and ISL1 through specific combinatorial regulatory logic. The bar plot (**top right**) illustrates prediction accuracy for each CR rule, with the x-axis indicating the index of CR. Line plots (left of the second and third rows) depict expression dynamics of FGF8 and GATA4 over *in silico* perturbation iterations under different configurations of the regulators, corresponding to CR1 and CR3. Boolean tables (bottom right) detail the groundtruth combinatorial logic for each rule. g. Same as in panel **f** but for the hematopoiesis system (Krumsiek et al. 2011).

After implementing the regulation prediction framework in Tabula, we apply it on four well validated regulatory networks involving hematopoiesis, cardiogenesis, neurogenesis and pancreatic endogenesis by zero-shot prediction without any fine-tuning. The core hematopoiesis network, one of the most studied systems, involves a total of 11 key transcription factors and 29 edges, which specifies cell fates of seven hematopoietic lineages (**Supplementary Fig. 7a and c**). The core cariogenesis network involves nine key TFs and signaling factors (e.g. BMP2, FGF8), and 16 edges, which controls the cell fate for the first heart field and second heart field (**Supplementary Fig. 7b and d**). The core neurogenesis network involves only seven transcription factors and a total of nine edges, which dedicates six neural lineages (**Supplementary Fig. 7e and f**). Lastly the core pancreatic endogenesis network includes eight TFs, 22 edges and involves eight pancreatic lineages across three time points at embryonic day 12 or E12, E14 and E17 (**Supplementary Fig. 8**). Surprisingly, Tabula’s simple in silico perturbation approach achieves an overall high predictive accuracy of > 0.83 across four systems on pairwise regulation prediction without the need for any fine-tuning, leveraging the same pretrained model. Taking the three-gene motif of *GATA1, FLI1*, and *KLF1* within the core hematopoiesis network (Qiu et al. 2022)(Krumsiek et al. 2011)(Qiu et al. 2022) (**Fig. 3c and d**) as an example, Tabula accurately predicts both the expression dynamics over iterations (**Fig. 3c**) and distribution shifts pre- and post-perturbation on either activation or repression regulations (**Fig. 3d**). Furthermore, Tabula accurately recovers 24 out of 29 pairwise interaction pairs by either expression distribution shifts (by default) or cross-cell regulation proportions (**Fig. 3e**). Moreover, the wrongly predicted perturbation pairs often involve genes with significantly lower gene expression compared to other genes (data not shown), underscoring Tabula’s reliability in determining regulatory modes.

It has long been known that the large cell type diversity results from the innumerable combinatorial gene regulations of a limited set of genes. Although massively parallel genetic screening, such as Perturb-seq, have made great strides in studying genetic interactions, the exponential increase in the possible higher order interactions render the effort in comprehensively profile combinatorial regulation an impossible mission. It is thus a great contribution if we are able to demonstrate foundation models are capable of revealing higher-order interactions. We thus take on the challenge to demonstrate Tabula’s ability to predict combinatorial regulation (CR) from the hematopoiesis system and the cardiogenesis system, both have very well validated combinatorial regulation. Overall, for each system, Tabula’s capability in revealing combinatorial rules (CRs) is evaluated against all boolean logic gates. For the cardiogenesis system, there are four known CRs. Our analysis demonstrated that Tabula achieved 100% consistency with the ground truth for CR1 and CR2, 87.5% for CR3, and 12.5% for CR4 (**Fig. 3f**) (**Supplementary Fig. 9a**). We next demonstrate the predicted dynamics of target gene expression for all possible regulator states across all in silico perturbation iterations. For example, FGF8 (CR1) shows two distinct trajectories for all combinations of regulator states (e.g., 0-0-0, 0-1-0, etc.), with the predicted expression levels distribution before and after in silico perturbation 100% aligning with the ground truth provided in the Boolean tables. For CR3, while the majority of predicted expression dynamics follow the same trend and align with the Boolean tables, one regulator state (0-0-0) shows a deviation and fails to capture the dynamics. For the hematopoiesis system, there are nine CRs (C. Wang et al. 2023)(J. Wang et al. 2020)(C. Wang et al. 2023). Tabula’s predictions achieve 100% consistency for CR1, CR6, CR8, and CR9, while CR3 and CR4 reached 81.25% and 75% consistency, respectively (**Fig. 3g**) (**Supplementary Fig. 9b**).

Taken together, we have demonstrated Tabula’s ability to accurately reveal pairwise and combinatorial gene regulation, generalizable across a large range of diverse biological systems, establishing one of the first testament in gaining mechanistic and predictive insight for a single cell foundation model.

## DISCUSSION

Advances in genomics now enable us to profile single cells at unprecedented scale, resolution and diversity across the spectrum of human biology. The expanding single cell dataset provides increasingly rich and diverse data, allowing the creation of virtual cell models to enhance our understanding of human biology with generative AI. Early efforts in this direction, such as Geneformer, scGPT and scFoundation, have demonstrated great promise. However, as we begin training foundation models on a large corpus of human data, data privacy becomes a significant concern. Moreover, while it is convenient to construct gene orders across cells to mimic word orders in sentences for subsequent transformer-based pretraining (as employed in these efforts), the unique features of single cell data demand novel design.

We developed Tabula, the first single cell foundation model that explicitly models the tabular structure of scRNA-seq and integrates federated learning to address privacy concerns. Tabula is unique in two key ways. First, it uses a tabular learning framework to explicitly model the tabular structure of single cell data. Unlike other approaches that preprocess gene expression into gene sequences, Tabula represents each cell as a permutation-invariant row of genes. Furthermore, rather than relying on mask language modeling, Tabular’s pretraining comprises of two objectives, namely gene-wise learning, which reconstruct original gene expression from a corrupted view, and cell-wise learning, which uses contrastive learning in the latent space to distinguish between positive (cell pairs from the same origin, with one generated from corrupted data) and negative (cell from distinct origins) cell pairs. Extensive benchmarking demonstrates that tabular learning outperforms MLM- and autoregression-based approaches on a range of cell- and gene-level downstream tasks, e.g. cell type annotation, batch correction, gene imputation and gene perturbation. Second, Tabula employs federated learning (Kairouz et al. 2021), allowing decentralized clients to train independently without sharing raw data, thereby addressing the privacy concerns. Thanks to the architectural improvement, we show that Tabular is able to allow impressive results in recovering pairwise and higher-order regulation across diverse biological systems. This work establishes one of the first pieces of evidence for the single cell foundation model successfully recovering ground-truth regulatory networks.

Although Tabula represents an important advancement in single cell foundation models, it is not without limitations. In its federated learning framework, all clients share a general transformer model, while distinct embedders are used for client-specific data. This design enables handling heterogeneous data distributions across clients (e.g., tissue-specific data). Ideally, however, a universal embedder could be used. Nevertheless, the client-specific embedder primarily affects zero-shot learning on downstream tasks, where tissue-specific embedders improve prediction performance. For fine-tuning downstream tasks, the choice of embedder has minimal impact. Currently, Tabula has been trained on only 15 million single cells, focusing solely on transcriptomics data. Scaling up training to include more single cells and incorporating multi-omics data, e.g. chromatin accessibility and gene expression co-assays, is an important next step. For instance, ideas from GET (Fu et al. 2024), a foundation model of transcription based on chromatin accessibility and sequencing data, could help extend Tabula from a single-modality foundation model to multi-modal foundation model. Although Tabula has shown promise in gaining mechanistic biological insights, such as high-order perturbation predictions, future work must further expand the prediction capabilities to reveal novel biological insights, such as the optimal gene combination that converts one cell fate to another. While single cell foundation models excel in various tasks, integration with more specialized models may be complementary in the short time.

Finally, with the rapid advancements of AI agents in recent years, combining foundation models with agent systems to automate biological discovery represents an exciting frontier in single cell biology (Xiao et al. 2024; B. Li et al. 2024). This endeavor could pave the way not only for virtual cells (Bunne et al. 2024) but also for virtual labs (Swanson et al. 2024) or even virtual scientists (Lu et al. 2024) in years to come.

## ACKNOWLEDGEMENTS

We appreciate valuable brainstorming, technical discussion, feedback from Y.M., W.T., and K.P. and other members in the Qiu lab (Stanford) and DSE lab (MSU) during the preparation and execution of this work as well as the writing of the manuscript.

## AUTHOR CONTRIBUTIONS

J.D., and Y.W. conceived the study, integrating tabular learning for single-cell data modeling with federated learning to ensure data privacy protection. J.D. led the development of the entire project. J.L and S.J contributed to the model preatraining and fine-tuned downstream task analysis with guidance from J.D., X.Q., M.L, and J.T. J.D. designed and generated the initial version of all figures in the manuscript with the help of J.L., S.J., and Z.M.; X.Q. provided additional guidance on refining the revised figures, from initial concept design to final organization. J.D., J.L., S.J., and X.Q. wrote the manuscript with feedback from all other authors. All authors read and approved the final manuscript.

## CODE AVAILABILITY

**Tabula** (version: 1.0) is implemented with Pytorch Lightning and freely accessible at https://github.com/aristoteleo/tabula.

## DATA AVAILABILITY

All data utilized in this study are publicly accessible, with detailed usage descriptions provided in the Methods section. The data sources are as follows: The pretraining datasets were obtained from the CELLxGENE census (https://cellxgene.cziscience.com). For the imputation task, both PBMC5K and Jurkat datasets, generated by 10x Genomics, are available from their official website (https://www.10xgenomics.com/datasets). The Melanoma dataset is publicly accessible in the GEO database under accession number GSE99330. For the genetic perturbation prediction task, Norman and Adamson datasets can be accessed via (https://dataverse.harvard.edu/api/access/datafile/6154020) and (https://dataverse.harvard.edu/api/access/datafile/6154417), respectively. The Replogle dataset was retrieved from (https://gwps.wi.mit.edu). For the cell type annotation task, the Human Pancreas dataset is available at (https://figshare.com/projects/TOSICA_demo/158489). The Myeloid dataset can be accessed from (https://drive.google.com/drive/folders/1VbpApQufZq8efFGakW3y8QDDpY9MBoDS). The 293t & Jurkat cells and DC datasets are available at (https://hub.docker.com/r/jinmiaochenlab/batch-effect-removal-benchmarking). The corresponding GEO database accession numbers for the liver reference and query datasets are GSE115469 and GSE124395, respectively. For the multi-omics integration task, the 10x Multiome PBMC dataset was obtained from (https://scglue.readthedocs.io/en/latest/data.html). The BMMC dataset is available in the GEO database under accession number GSE194122. The Perirhinal Cortex dataset can be accessed through (https://cellxgene.cziscience.com/collections/283d65eb-dd53-496d-adb7-7570c7caa443). For the in silico gene regulatory network inference task: the hematopoiesis dataset is available at (https://dynamo-release.readthedocs.io/en/latest/notebooks/tutorial_hsc_velocity.html); the cardiogenesis dataset is accessible in the GEO database under accession number GSE126128; the neurogenesis dataset can be retrieved from (mousebrain.org); and the pancreatic endogenous dataset is available in the GEO database under accession number GSE101099.

## ONLINE METHODS

### Pretraining data collection and preprocessing

#### Data collection

To establish a robust foundation for Tabula self-supervised pretraining, we assemble an extensive single cell RNA sequencing (scRNA-seq) dataset comprising transcriptomic data from approximately 15 million human cells, sourced from the CELLxGENE portal (https://cellxgene.cziscience.com/).

Given that the CELLxGENE portal is regularly updated, we utilized the release from July 25, 2023. The dataset encompasses cells from a range of major human tissues, including the intestine, pancreas, lung, heart, blood, kidney, and brain. Furthermore, cells from a variety of less-represented human tissues like the spinal cord and spleen were aggregated into a separate category called “others”. The compilation resulted in a total of 8 categories, namely “intestine”, “pancreas”, “lung”, “heart”, “blood”, “kidney”, “brain” and “others”, comprising 37.99 million human cells. In our federated learning framework, we set the number of clients to be eight, each corresponding to a distinct tissue category aforementioned in our setting. To maintain data balance across categories and guarantee effective pretraining, a selection process was implemented, limiting each category to a maximum of 3 million cells.

We regrouped each category according to its original dataset identifiers. We then performed quality control and data preprocessing procedures for cells belonging to the same source dataset separately (detailed methodology described in the next subsection). Within each category, we randomly selected 3 million cells from the preprocessed datasets. For categories with fewer than 3 million total available cells, all cells were used. This process ultimately yields a pretraining dataset of 15 million cells. The number of pretraining cells for each tissue client is as follows: Intestine: 80,060 cells; Pancreas: 220,436 cells; Lung: 3,013,840 cells ; Heart: 1,768,184 cells; Blood: 3,057,298 cells; Kidney: 816,538 cells ; Brain: 3,024,233 cells; Others: 3,046,184 cells.

#### Quality control & data preprocessing for federated learning

We implemented a dataset-specific strategy for quality control and data preprocessing. This strategy diverges from previous approaches of consolidating data prior to quality control and data preprocessing, allowing us to preserve the distinctive characteristics of each original dataset and reduce the impact of the batch effect. Specifically, we filtered cells with fewer than 250 detected genes to exclude potential empty droplets, non-viable cells, or severely degraded cells. Concurrently, we also removed genes detected in fewer than 250 cells to mitigate the influence of lowly expressed or rare transcripts. To capture biologically relevant variations, we identified 1,200 highly variable genes (HVGs) within each dataset after quality control. It is worth noting that HVGs were identified prior to the aggregation of datasets for each tissue. Consequently, HVGs may differ among datasets even within the same tissue. We chose this strategy because selecting HVGs from individual datasets or distinct contexts can improve the model’s ability to discern cell-specific characteristics while also enhances the model’s capacity to learn generalizable features from varied biological signals. To standardize the data for pretraining and further mitigate batch effects, we implemented a value binning technique, discretizing the expression values into 50 bins for each cell. This transformation normalizes the scale of gene expression across batches, creating a more uniform input space for the pretraining process. The choice of 50 bins strikes a balance between preserving gene expression differences and reducing batch-specific technical variations.

#### Data preprocessing for tabular modeling

Our model employs a tabular learning approach for self-supervised pretraining, with learning objectives that include contrastive loss and reconstruction loss. The contrastive loss focuses on enhancing feature discrimination between cells, while the reconstruction loss aims to capture complex relationships among genes within each cell. To achieve these objectives, we need to corrupt the input samples to construct corrupted views from the original views. We followed Xtab (Zhu et al. 2023) to construct corrupted views through random feature resampling. Specifically, we randomly selected a subset of genes in cells in a training batch and then resampled their values from the empirical marginal distribution (“Marginal Distribution” 2004) of these genes within the same training batch. We set the corrupted ratio at 60%. This means that for each cell sample’s original view *x* and its corrupted view 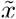, 60% of the gene values are resampled while 40% remain unchanged. This resampling strategy will provide sufficient perturbation to enhance the model’s robustness while retaining enough original information to maintain biological relevance.

### Tabula model architecture

Tabula is a privacy-preserving foundational model for single cell omics developed within a federated learning framework. Tabula also innovatively treats single cell data (represented as a cell-by-gene matrix) similarly to tabular data and incorporates a tabular transformer with attention mechanisms to facilitate the unsupervised learning of gene and cell embeddings. Additionally, Tabula handles cross-tissue variations by partitioning the models into different clients with tissue-specific embedders and project heads that capture the characteristics of each tissue, and a shared tabular transformer that stores the common knowledge across tissues. Note that this cross-tissue specific client system is implemented to demonstrate the federated learning, however, in general federated learning can be used for cross-institute learning to address privacy concerns by avoiding data sharing across institutes while benefiting the knowledge learned from the data produced from other institutes during pretraining.

In the following, we will describe the four main modules related to the Tabula model architecture, namely the embedder module, the tabular transformer module, the pretraining objectives and the collaborative federated learning framework. The embedder module transforms the original cell-by-gene table into embeddings, which are then used as inputs for the tabular transformer. Then the tabular transformer, powered with FlashAttention-2 (Dao 2023) takes the embeddings to learn contextual gene and cell representation efficiently and effectively. The output of the tabular transformer is further processed by project heads to derive the pretraining losses. Tabular pretraining objective is based on self-supervised loss functions tailored for cell-by-gene table learning through cell-wise row learning and gene-wise column learning. the collaborative federated learning framework module details the process of updating between the local and global tabular transformer models.

#### Embedder module

scRNA-seq data can be represented as a cell-by-gene matrix, **X** ∈ ℝ^*N* ×*M*^, where each entry **X**_*i,j*_ ∈ ℝ^*+*^ denotes the read count of RNA molecules for gene *j* ∈ {0,1, …,*M*}in cell *i* ∈ {0,1,…,*N*}. Each row of the cell-by-gene table, **X**, is considered as an input cell sample in a tabular transformer, and each column is a gene feature token. The function of the embedder module is to convert each cell sample to feature embeddings **E** ∈ ℝ^(*M+*1) ×*d*^. Here, *M* denotes the number of columns, +1 indicates the special [CLS] token (see below) and *d* is the embedding dimension. Tabula processes each row of the table as an unordered sequence of genes, where gene in each column is represented by its column feature name embedding (gene token embedding) and its corresponding feature value embedding (gene expression value embedding). It is important to note that even though the cell-by-gene table we are encoding in Tabula solely consists of gene tokens and does not contain metadata column features, such as cell type, tissue type, or sequencing technology, it is entirely feasible to encode metadata as well and we leave it for future work.

- **Column feature name embedding**. The feature column in the cell-by-gene table for each client corresponds to individual genes. Our pretraining dataset includes 196 distinct studies, from which we select 1,200 HVGs per study, yielding a total of 23,156 HVGs across all datasets. Consequently, we have 23,156 unique feature columns to learn during the pretraining phase. We assign each column feature name, *feature*_*j*_, a unique integer identifier, *id* (*feature*_*j*_) These identifiers collectively form the column feature name vocabulary utilized in Tabula. Additionally, we include a special token, [CLS], in the vocabulary, which is used to aggregate all column features into a cell-level representation. The input column feature tokens for cell *i* are therefore represented as a vector 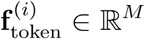, where *M* is a predetermined maximum sequence length in the tabular transformer:

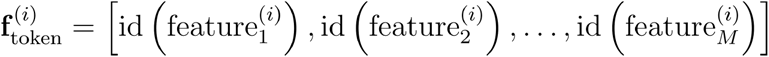

We then utilize the standard Embedding layer, which functions as a lookup table that maps discrete indices (e.g., unique id of column feature names) to continuous dense vectors, or embeddings. The embedding representation of input column feature tokens for cell *i* is denoted as 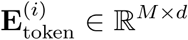 where *d* represents the embedding dimension:

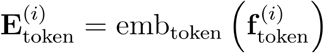

where emb_token_ represents an embedding layer to learn feature token embeddings.
- **Column feature value embedding**. Different sequencing technologies, with variable sensitivity, sequencing depth, and ability to detect lowly expressed genes, resulting in different data distributions and scales, introduce complexities into scRNA-seq data modeling (Sarkar and Stephens 2021). For instance, some platforms capture more genes but at lower depth, while others focus on a subset of genes at higher resolution (Svensson et al. 2017). These differences lead to inconsistent gene expression measurements, making it challenging to integrate data from diverse platforms for accurate modeling. To tackle this issue, we adopt scGPT’s value binning technique (Cui et al. 2024) to convert expression counts into relative values. Unlike scGPT, which only focuses on non-zero expression counts in each cell, we select 1,200 HVGs for modeling in each dataset, including those with zero expression. By incorporating zero-expressed genes, our method Tabula provides a more complete view of gene expression variability, capturing both active and inactive genes. This broader perspective allows for the detection of important biological patterns that could be overlooked when considering only non-zero genes, as zero expression can have biological significance, such as in regulatory processes (Jiang et al. 2022). For each HVG expression count in a cell, we compute the raw absolute values and partition them into *B* consecutive intervals [*Bin*_*t*_, *Bin*_*t*+1_], where *t* ∈ {1,2, …., *B*} The binned expression value 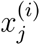 for cell is designated as:

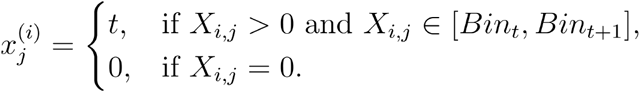

Of note, for certain fine-tuning tasks, if we initially select more than 1,200 HVGs, we also filter out zero-expressed genes from within those HVGs. The column feature values for cell *i* are therefore represented as a vector 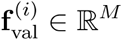, where *M* is a predefined maximum sequence length in the tabular transformer:

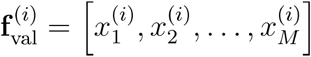

The embedding representation of input column feature values for cell is denoted as:

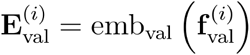

where 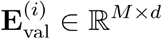, and emb_val_ represents an embedding layer to learn feature value embeddings.

Consequently, the embedding **E**^(*i*)^ ∈ ℝ ^*M* × *d*^ for cell *i* is represented as:

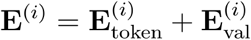

Finally, the embedding of the [CLS] token is appended to the feature embedding, resulting in a final feature embedding **E**^(*i*)^ ∈ ℝ^(*M*+1)×*d*^. The final hidden state corresponding to the [CLS] token is used as row (cell) representation aggregating from column feature embeddings. It is important to note that each client’s embedder module is distinct and specialized for learning tissue-specific features.

#### Tabular transformer

The backbone of the tabular transformer is based on self-attention transformer (Vaswani et al. 2017). The local tabular transformers and global tabular transformers share the same structure among clients. The tabular transformer on each client is designed to learn specific knowledge for different tissues during the pretraining stage. Our tabular transformer treats each gene column as an input token and aims to capture column gene dependency for each cell. The tabular transformer takes **E**^(*i*)^ ∈ ℝ^(*M*+1)×*d*^ as input a set of *M* gene column embeddings along with a [CLS] token embedding. The input to the first layer of the stacked tabular transformer is denoted as 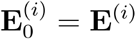. The output of the stacked transformer blocks is defined as follows:

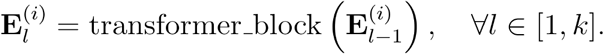

where *l* represents the number of stacked transformer layers. The core function in transformer_block is the attention mechanism that can be formulated as:

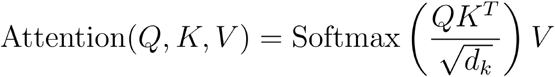

where *Q* = *EW*_*q*_, *K* = *EW*_*k*_, and *V* = *EW*_*v*_ denote linear transformations of the input *E*. Specifically, *Q, K*, and *V* correspond to the query matrix, key matrix, and value matrix, respectively. *W*_*q*_, *W*_*k*_, and *W*_*v*_ are training parameters. *d*_*k*_ is the dimension of the key vectors. The softmax function is applied to the scaled dot-products of the queries and keys to produce attention weights, which are then used to weigh the values. This mechanism allows the model to focus on different column genes of the cell input dynamically, enabling it to capture complex gene dependencies effectively. To optimize the efficiency of self-attention mechanisms, we employ FlashAttention-2 (Dao 2023) accelerated implementation. This advanced approach enhances its ability to handle large input dimension *M* effectively. It is important to note that the current transformer in Tabula can be replaced with any other efficient transformer.

The output 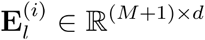 incorporates updated *M* gene embeddings and a [CLS] embedding, which represents the cell embedding. This representation is automatically aggregated from all column gene embeddings through an attention mechanism. For pretraining and fine-tuning, the transformer output is processed by a projection head, typically a Multilayer Perceptron (MLP) (Taud and Mas 2018). This MLP handles the embeddings from the transformer, facilitating pretraining tasks like predicting masked gene columns and fine-tuning for specific tasks such as cell type classification. The [CLS] embedding can then be extracted to evaluate and visualize the effectiveness of the learned cell representations.

#### Tabular pretraining objective

Current foundatiol models for single cell analysis draw parallels between single cells and natural language, where texts are composed of words, and cells are defined by genes. Whether employing autoregressive self-supervised learning as in scGPT (Cui et al. 2024) or mask-reconstruction pretraining used in Geneformer (Theodoris et al. 2023), scFoundation (Hao et al. 2024), and scBERT (Yang et al. 2022), these methods treat scRNA-seq data as ordered gene sequences. They either use attention maps to artificially impose an order, or rank genes to create ordered sequences. Both modeling approaches contrast with the inherent unordered nature of single cell omics. Additionally, natural language consists solely of discrete word token IDs, whereas scRNA-seq data includes both discrete gene token IDs and continuous numerical gene values. Instead, we represent scRNA-seq data as tabular data, with columns corresponding to gene features and rows representing cells.

Swapping any two gene columns does not alter the properties of the cells. Feature values can be either numerical or categorical; however, this work focuses exclusively on numerical values of genes while leaving metadata information for future study. To effectively model the cell-by-gene table, we design two specialized pretraining losses for tabular learning: reconstruction loss and contrastive learning to learn gene column features and row cell features respectively.

To calculate the reconstruction and contrastive losses, a corrupted view must be generated for each cell. The corrupted view is created through random feature resampling. For example, for cell *i*’s original view **c**^(*i*)^ ∈ ℝ^*M*^ where *M* represents the number of gene columns and the gene values in **c**^(*i*)^ have been already binned, we randomly choose a subset of *M* genes from cell *i* as genes whose expression will be corrupted and resample their values from the empirical marginal distribution (“Marginal Distribution” 2004) of these genes within the training batch. We set the corrupted ratio at 60%. It is worth noting that the corrupted genes for each cell differ from one another. The corrupted cell-wise view for cell *i* is denoted as 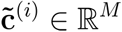.

- **Reconstruction loss as gene-wise column learning**. Reconstruction loss is a self-supervised training objective. It aims to recover the original gene-wise view from a corrupted view of that gene. Taking gene *j* as an example, we take the corrupted view, 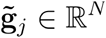, for gene *j* as input and aim to reconstruct its original binned gene expression **g**^(*i*)^ ∈ ℝ^*N*^ where 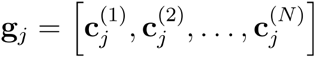 The reconstructed view for gene *j* can be represented as 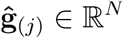 In this study, we use Mean Squared Error (MSE) to measure the distance between the original view and the reconstructed view. The reconstruction loss is denoted as:

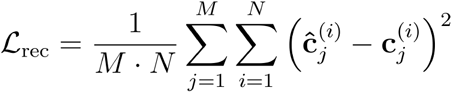

It is crucial to highlight that the reconstruction loss ℒ_rec_ focuses on reconstructing all *M* gene values at the column (gene) level, not just the corrupted ones. By training the model to restore gene expressions, the model develops a deep understanding of each gene’s distribution and its relationship with other genes. This leads to more accurate gene-specific feature representations. It is important to highlight that existing foundation models (Theodoris et al. 2023; Hao et al. 2024; Yang et al. 2022) typically assign a mask or corrupted value of -1 or 0, which does not align with the conditions encountered during the inference stage, where the gene values are not set to -1 or 0. Instead, we replace the corrupted values by resampling from the empirical marginal distribution of all corrupted genes, ensuring consistency with the inference process.
- **Contrastive loss as cell-wise row learning**. Similar to the reconstruction objective, we generate 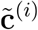 as a corrupted cell-wise view of the cell *i*. **c**^(*i*)^ and its corresponding corrupted view 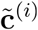 are treated as a positive cell **c**^(*i*)^ pair, while and other cells **c**^(*i*)^ within the batch are treated as negative cell pairs. In general, contrastive loss aims to minimize the distance between positive pairs of sample cells and maximize the distance for negative pairs. In this study, we employ the SimCLR (T. Chen et al. 2020) loss for contrastive pretraining, which is denoted as :

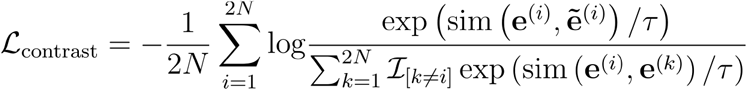

where **e**^(*i*)^ and 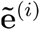 represent the cell embedding of cell *i* from the original view and corrupted view respectively after being processed through the transformer block. ℐ_[*k*≠*i*]_ ∈{0,1}is an indicator function that takes the value of 1 if *k*≠*i*; otherwise, it is 0. The similarity function *sim* (·, ·)represents cosine similarity here to measure the similarity between two cell embeddings, τ denotes the temperature parameter and *N* is the number of cells in a batch.

The final pretraining objective in Tabula is a combination of the two loss functions:

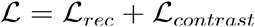

The overall loss combining reconstruction and contrastive losses offers a holistic approach to learning both gene-level (column) and cell-level (row) features from the cell-by-gene table.

#### Collaborative federated learning framework

Tabula is based on the federated learning framework. Unlike existing foundation models based on centralized learning, which require aggregating all data in a central location and thus raise privacy and ethical concerns, Tabula allows data to remain distributed across multiple clients. Each client trains a local model on its own client data, and only model parameters are shared with the central server for global model updates. To be specific, the key steps are as follows:

- **Local training**. Each client *k* (e.g., institution or hospital) trains a local copy of the model using its own data *D*_*k*_. During this step, the model weights *W*_*k*_ are updated by minimizing the local loss function ℒ_*k*_ (*W*)based on the client’s local dataset. The local training process can be denoted as:

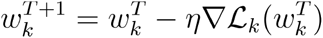

where *T* is the current iteration, *η* is the learning rate and 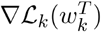 is the gradient of the loss function at the current iteration weights 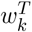. The model weights contain two components: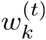 is the model weights for tabular transformer and 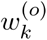 is the modelweights for the rest of architecture including embedders and project heads. 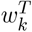 can be represented as 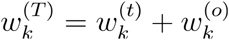.
- **Sending local updates**. Once local training is complete, each client *k* sends its updated tabular transformer gradient 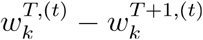 to a central server. These gradients represent the local knowledge learned from the individual datasets. This can be summarized as:

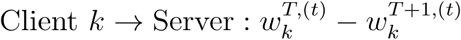
- **Global aggregation**. The central server aggregates the local tabular transformer updates from all clients to create a new global tabular transformer. In Tabula, we adopt a common aggregation method which is Federated Averaging (FedAvg) (X. Li et al. 2019). The global tabular transformer weights 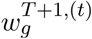 are updated by computing a weighted average of the local tabular transformer updates, where the gradients reflect the significance of each client. The importance of each client is quantified by a corresponding weight *P*_*k*_. It can be summarized as:

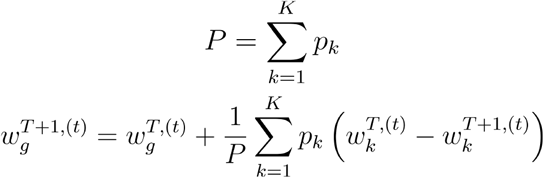
- **Broadcast global update**. After aggregation, the global tabular transformer is updated with the newly averaged weight 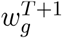.This updated global model is then broadcasted back to the clients, which subsequently update their local models based on the received global weights. This process can be expressed as:

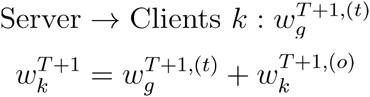

Those processes are repeated for several rounds, with clients continuing to train the model locally, send updates, and the server performing global aggregation, until the model converges or reaches the desired performance. In Tabula, we update the global model once every epoch and treat each client with the same importance weight.

Our federated pretraining framework offers three key advantages. First, it ensures data privacy, as sensitive information remains localized on the client side, preventing unnecessary data transfers. Second, it distributes the computational burden across multiple clients, alleviating the demand for resources on any single entity and enhancing scalability. Third, the federated architecture naturally enables the development of tissue-specific embedders and project heads, which are designed to capture tissue-specific features. In contrast, conventional foundation models rely on a single, generalized embedder, making them less effective at capturing the nuances of tissue-specific variations. It is important to note that in our federated training framework, both the local model and the tissue-specific embedder, along with the project head, can be shared. This allows for greater flexibility and collaboration across clients while preserving the benefits of tissue-specific feature learning.

#### Similarity and innovation compared to existing foundation models

Compared to existing foundation models for single cell data, Tabula presents four key innovations. First, Tabula utilizes federated learning instead of centralized training, ensuring enhanced data privacy by allowing institutions to keep sensitive data local while only sharing model updates. Second, Tabula introduces an innovative approach by treating single cell data as tabular data and designing two distinct pretraining losses specifically for tabular modeling. Reconstruction loss is designed for column-level learning to capture gene-related features, while contrastive loss focuses on row-level learning to highlight distinctions among individual cells. This dual mechanism allows the model to have a holistic view to understand tabular data. Third, we are the first to propose tissue-specific embedders and project heads in the field of foundation models for single cells. This novel approach allows the model to tailor its representations to the unique characteristics of each tissue type, enabling more precise feature extraction compared to traditional models that rely on a single, generalized embedder. Finally, Tabula is a lightweight foundation model, achieving comparable performance to larger models like scGPT with significantly fewer parameters, making it more efficient and scalable for real-world applications.

### Fine-tuning on downstream tasks and analysis

- Cell type annotation The performance of cell type classification was evaluated through metrics including accuracy, precision, recall, and F1 score (See **SUPPLEMENTARY NOTES S1**), offering a well-rounded view of Tabula’s predictive accuracy against other models. Additionally, the confusion matrix was built to reveal patterns of predictions across cell types. To fine-tune the pretrained Tabula for the cell type annotation task, an MLP classifier is introduced. This classifier takes cell embeddings generated by the Tabular Transformer as input and outputs categorical predictions for cell types. The entire model is optimized using cross-entropy loss ℒ_*AN*_, as depicted below:

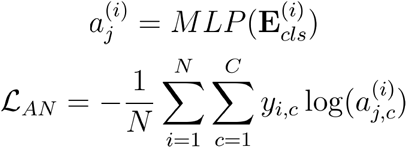

Here, the MLP layer outputs the predicted probabilities of cell type labels 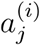 taking the cell representation 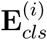 as input. The cross-entropy loss, ℒ_*AN*_, is computed where *N* denotes the total number of samples in a batch, *C* is the number of cell type classes, *y*_*i, c*_ represents the true label for *i*-th sample for class *c*, encoded in one-hot format, and 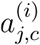 is the predicted probability for the *i* -th sample that belongs to cell type class *c*. We assessed our model using four datasets including Human pancreas (hPancreas), Myeloid, Cell Lines, and Liver datasets (See **Downstream Task Datasets**). Benchmark comparisons were made against scGPT, Geneformer, and scFoundation. The fine-tuning settings of these benchmark models follow their default settings.
- Multi-omics integration To assess Tabula’s performance in integrating cell embeddings of multi-omics data after fine-tuning, we employ biological conservation metrics (Malte D. Luecken et al. 2022) including Normalized Mutual Information (NMI_cell_), Adjusted Rand Index (ARI_cell_), and Average Silhouette Width (ASW_cell_) (See **SUPPLEMENTARY NOTES S2**). These metrics quantify the alignment between cluster formations derived from the embeddings and the validated ground truth cell type annotations. Consistent with methodologies used in previous benchmarking studies (Cui et al. 2024), we calculated the average of these three metrics (AvgBIO) to provide a composite score reflecting overall performance in the task. During the fine-tuning phase, the model weights are refined using multiple loss functions to enhance the learning of aligning different omics modalities. Specifically, the Tabula employs three fine-tuning losses: the reconstruction loss ℒ_rec_ from the pretraining stage (See **Tabular pretraining objective**), masked gene modeling (MGM) by applying masking to random-selected genes, and cell context-aware masked gene modeling (CMGM), which integrates cell-level information, as detailed below:
  - Masked gene modeling To elucidate gene interrelationships, Tabula employs MGM loss as a fine-tuning objective. This approach parallels the reconstruction loss employed during the pretraining but modifies the handling of genes at these masked positions. Specifically, whereas the reconstruction loss utilizes a corrupted view for masking, MGM involves the random masking of a subset of gene expression values *e*^*i*^ in each input cell. Tabula is then optimized to predict these masked expression values accurately. This setup enables the model to infer gene expression patterns from other genes within cells, thereby enhancing inter-gene awareness. The training objective is quantified using the mean squared error (MSE) at the masked positions, denoted as ℳ_mask_. The formula is depicted below:

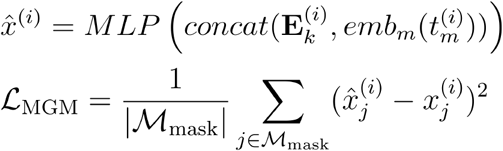

Here, 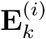 denotes the gene embeddings from the Tabular Transformer layers. By concatenating the embedding of modality information 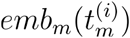 generated by a modality encoder given a modality label 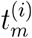, the integrated embedding will be passed to an MLP to predict the gene expression value 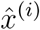 for cell *i*.
  - Cell context-aware masked gene modeling Following the previous work (Cui et al. 2024), the CMGM loss advances model optimization by predicting gene expression values through incorporating cell representation, which promotes contextual awareness between cells and genes and enhances the efficacy of cell representation learning. Specifically, a gene-specific query vector *q*_*j*_ is created given the gene token embedding 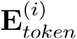 and calculates the predicted expression value using the parameterized inner product between *q*_*j*_ and the cell representation 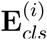:

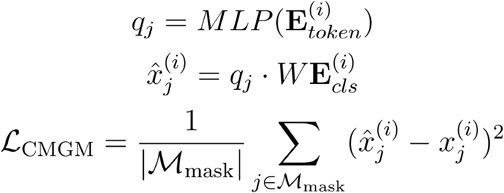

Here, 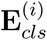 denotes the cell-level embedding extracted from the [CLS] token. ℒ_CMGM_ is computed as MSE loss between the predicted expression value and true value, as similar to ℒ_*MGM*_.
- Multi-batch integration To access modeling performance on multi-batch integration, biological conservation metrics (Malte D. Luecken et al. 2022), especially the inverse of the average silhouette width for batch clustering (*ASW*_*batch*_) and graph connectivity (*Graph_Conn*) are employed. The overall score, *Avg*_*batch*_ is computed by averaging the two metrics (See **SUPPLEMENTARY NOTES S2**). During the fine-tuning phase, a domain adaptation (DA) loss is utilized to enable the model to optimize batch correction by reversing the backpropagation of learned batch labels. Following the methodology used by scGPT, we initiate an MLP classifier that predicts the batch label based on the cell representation 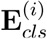. Through a reverse back-propagation procedure, the model will aggregate batch information by integrating the reversed gradients, thereby enhancing the robustness of the batch correction process. Cross-entropy loss is employed for model finetuning. The formula is illustrated as follows:

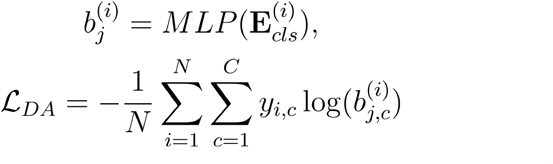

Here, the MLP layer outputs the predicted probabilities of batch labels 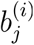 using the cell representation 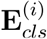 as the input. The domain adaptation loss, ℒ_*D,A*_, is computed as the negative categorical cross-entropy loss, where *N* is the total number of samples in a batch, *C* is the total number of batch *y*_*i, c*_ classes, is the true label for *i*-th sample for class *c*, in one-hot encoded form, and 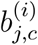 is the predicted probability for the *i* -th sample belongs to batch class *c*. We conducted a comprehensive evaluation of multi-omics and multi-batch integration tasks within a unified framework, with four losses: ℒ_rec_, ℒ_*MGM*_, ℒ_*CMGM*_, and ℒ_*DA*_, optimized together according to the batch and omic features in the dataset. For these evaluations, we use datasets including 10x Multiome PBMC, BMMC, Perirhinal Cortex, Human dendritic cells (See **Downstream Task Datasets**). Benchmark comparisons were made against scGPT (Cui et al. 2024), Geneformer (Theodoris et al. 2023), and scBERT (Yang et al. 2022). The fine-tuning protocols for these benchmarking models adhered to the methods described for scGPT.
- Genetic perturbation prediction We assessed the performance on genetic perturbation prediction using two key metrics: Pearson_*delta*_ that quantifies the correlation between predicted and actual expression changes after perturbation, and DEG Pearson_*delta*_ that specifically evaluates this correlation for the top 20 most differentially expressed genes (See **SUPPLEMENTARY NOTES S3**). During the model testing, there are four distinct scenarios pertaining to gene perturbation visibility as defined in GEARS (Roohani, Huang, and Leskovec 2023): (i) 1/1 Unseen: single-gene perturbations are not previously observed during the model training; (ii) 2/2 Unseen: two-gene perturbations that neither gene are not previously observed during the model training; (iii) 1/2 Unseen: two-gene perturbations where one gene is previously unobserved during the model training; and (iv) 0/2 Unseen: two-gene perturbations with both genes having been previously observed during the model training. These evaluations help elucidate the model’s predictive robustness across varying levels of experimental uncertainty. During the fine-tuning stage, a perturbation encoder (MLP) is initialized for incorporating perturbation information through adding a perturbation embedding 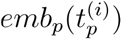 to the original input. Then, 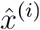 is computed by passing the integrated embedding to the transformer layers and a reconstruction head. The ℒ_*pert*_ is calculated as MSE loss across all the positions. The process is outlined in the formula below:

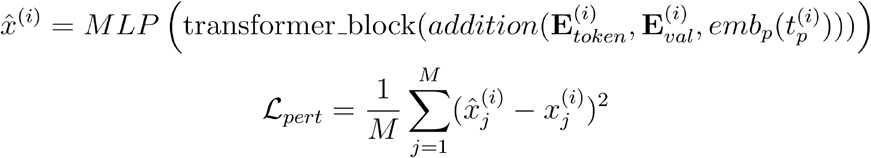

Here, 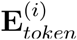 and 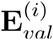 are representation embeddings of gene token and gene expression value, respectively. We evaluated our model using three Perturb-seq datasets derived from leukemia cell lines, including Adamson, Norman, and Replogle (See **Downstream Task Datasets**). Adhering to preprocessing protocols established by the recent perturbation prediction methodology, GEARS (Roohani, Huang, and Leskovec 2023), we benchmarked Tabula against scGPT, scBERT, and Geneformer. This comparison was conducted in a consistent fine-tuning framework, utilizing gene-level embedding and perturbation information to predict perturbed expression profiles.
- Reverse perturbation prediction For the reverse perturbation prediction task, we aimed to identify perturbation conditions that best replicate observed gene expression profiles from a query set. We followed the setting of scGPT (Cui et al. 2024) to select a subset of 20 genes from the Norman dataset, which consisted of 39 cases for training, 3 for validation, and 7 for testing. These 20 genes include a perturbation space comprising 210 combinations, encompassing both single-gene and double-gene perturbations. For each test case, we treated the ground truth gene expression profiles under the test perturbations as queries and all 210 perturbation conditions as references for retrieval. Following the methodology outlined in scGPT, we formulated the retrieval process as a top-K retrieval task with a two-round selection strategy involving Euclidean distance-based ranking and ensemble voting to identify the most-likely perturbation conditions. To construct the reference database, we generated 30 distinct gene expression profiles for each of the 210 perturbation conditions by randomly sampling control cells, resulting in 6,300 post-perturbation profiles:

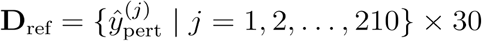

This approach allowed for increased diversity and robustness in representing each perturbation condition. For each test perturbation *X*+*Y*, the ground truth gene expression profiles from cells undergoing *X*+*Y* perturbation were treated as the query set:

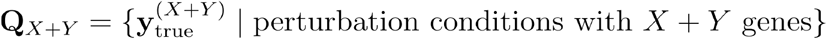

The retrieval strategy involved two rounds. In the first round, each cell within the query set, *Q*_X+Y_, identified its top K closest profiles from the reference database, *D*_ref_, by minimizing the Euclidean distance:

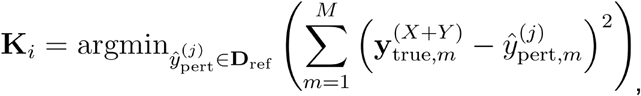

where *K*_*i*_ denotes the set of top K most similar expression profiles for each query cell from *D*_ref_. In the second round, we implemented an ensemble voting scheme: each query cell cast a vote for its top K matches within *D*_ref_. Perturbation conditions were then ranked by the total number of votes received from all cells in the query set:

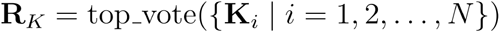

The conditions with the highest votes were identified as the predicted perturbation conditions, which we reported in descending order based on vote count. For evaluation, we employed a modified top-K accuracy metric to assess retrieval performance. This metric accounted for two criteria: (1) exact matches, where the predicted perturbation condition precisely matched the ground truth condition, and (2) partial matches, where at least one gene in the predicted condition overlapped with the actual perturbation combination:

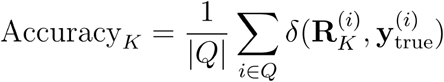

Here, *δ* denotes the evaluation function that returns a score of 1 for an exact or partial match, and *Q* represents the query set. To ensure robustness, we averaged retrieval performances across five random seeds. We benchmarked Tabula against scGPT on both exact match and partial match settings.
- Imputation We assessed the imputation performance using several quantitative metrics (See **SUPPLEMENTARY NOTES S4**). The primary evaluation involves the Pearson correlation coefficient (Pearson), Root Mean Squared Error (RMSE), and Mean Absolute Error (MAE) to measure the accuracy of imputed values compared to the true values. To demonstrate the model’s robustness, we applied different masking ratios (10%, 20% and 30%) to the dataset, with Pearson correlation as one of the metrics. Additionally, the model’s performance is analyzed through Mean Squared Error (MSE) at both the cell-wise and gene-wise levels. To assess the quality of imputation, cosine similarity is calculated between different cell types. This helps evaluate how well the imputed data preserves the original structure and relationships of the data. Specifically, it provides insights into the accuracy of the imputation and ensures that the natural diversity and variability between cell types remain consistent before and after the imputation process. Since the true dropout values are unknown, we evaluated the method’s performance by randomly masking the expression matrix in the scRNA-seq dataset, replacing these values with zeros. To more accurately simulate the distribution of dropout values, we adopted the settings from DeepImpute(Arisdakessian et al. 2019), and utilized an exponential kernel(Wen et al. 2023). For each cell, we identify genes with non-zero expression values, excluding cells with few expressed genes. For these non-zero genes, we compute sampling probabilities using an exponential distribution and normalize them to sum to 1. Using these probabilities as weights, we randomly sample a subset of the non-zero genes without replacement. The expression values of the selected genes are then masked to zero to simulate dropout genes, while the remaining genes retain their original expression values. The model is then used to reconstruct the true expression values of these masked genes. During the fine-tuning process, the masked data is divided into three sets: the training set, the validation set, and the test set. For each set, predictions are made exclusively for the positions within that specific set. Subsequently, an imputation decoder (MLP) is introduced for each gene, with the objective of predicting the expression levels at the respective positions. 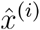 is computed by passing the integrated embedding to the transformer layers and imputation decoder. We calculate the imputation loss ℒ_*imp*_ in masked positions. The process is outlined in the formula below:

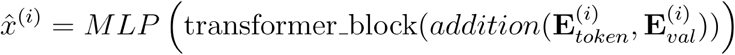

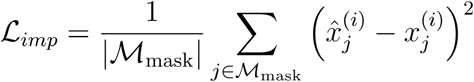

Here, 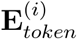 and 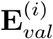 are representation embeddings of gene token and gene expression value, respectively. We assessed our model using four datasets derived including PBMC5K, Jurkat, Melanoma, and hPancreas (See **Downstream Task Datasets**). We benchmarked Tabula against scGPT, scBERT, and Geneformer. This comparison was conducted in a consistent fine-tuning framework, utilizing gene-level embeddings for the prediction of dropout expression profiles.
- In silico gene regulatory network inference For the in silico inference on gene regulatory networks, we performed an iterative process on our pretrained Tabula model to explore the impact of incremental increases in gene A expression on gene B. Initially, the expression levels for all genes in a given cell are captured, with specific focus on genes A and B, denoted as *y*_0_ and *x*_0_, respectively. At the initial conditions, gene expressions of a cell are illustrated as **G**_0_ = {*g*_1,0_, *g*_2,0_, …, *g*_*n*,0_}, where *g*_*i*,0_ represents the initial expression level of the *i*-th gene, with *g*_*A*,0_ = *y*_0_ and *g*_*B*,0_ = *x*_0_. Then, each iteration involves modifying the expression level of gene A by incrementing it by one, while maintaining the context of all other genes from the last iteration’s output: *g*_*A,t*+1_ = *g*_*A,t*_+1. The entire gene expression profile, including the updated gene A, is input into the Tabula model, the output is then obtained from the pretrained reconstruction head. The iterative process is illustrated below:

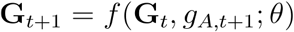

Here, **G**_*t*_ represents the gene expression profile at time *t, f* and is the function modeled by the pretrained Tabula, encapsulating the parameter θ. The continuous output expression value of gene B is recorded at each iteration step. This pseudo time-series data of gene B’s expression levels provides a quantitative measure of the dynamic response to the systematic perturbation of gene A. Let *X* = {*x*_1_, *x*_2_, …, *x*_*t*_} denotes the sequence of gene B’s expression levels over *T* iteration steps. Based on the continuous output sequence for gene B, one can plot the trajectory of gene B’s expression levels across the iteration steps. By observing the trend in these values, it is possible to infer the regulatory effect of gene A on gene B. Should the expression levels of gene B consistently increase, this suggests that gene A acts as an activator. Conversely, a decline in gene B’s expression levels would imply a suppressive influence by gene A. Building upon the in silico perturbation prediction of pair-wise gene regulation e, we extended our approach to combinatorial regulation predictions. In this more complicated scenario, we conducted in silico perturbations on multiple genes simultaneously and observed the continuous output expression of the target gene, following the logic established above. We accessed gene regulatory rules across four biological systems including hematopoiesis, pancreatic endogenesis, neurogenesis, and cardiogenesis (See **Downstream Task Datasets**). For the evaluation of combinatorial regulation logic, our analysis is concentrated on the hematopoiesis and cardiogenesis systems, the only two systems with very known combinatorial gene regulation logic.

### Implementation details

Tabula was implemented in Python and utilized PyTorch Lightning to construct its overall framework. Within the federated learning setup, message passing interface (MPI) was employed in the cluster to allocate computing nodes to each client. The NCCL framework (Nvidia 2017) was adopted for efficient communication among clients over the InfiniBand (IB) network. Additionally, torch.distributed was implemented to realize the message queues of the parameter sending, receiving, broadcasting and managing process.

- Tabula Tabula employs FlashAttention-2 (Dao 2023) architecture, comprising three transformer blocks each containing 8 attention heads, with an output embedding size of 192. The maximum input gene length was set to 1,200, corresponding to 1,200 highly variable genes (HVGs) selected from each dataset. The pretraining dataset consisted of 15 million cells, which were randomly sampled from tissue-specific subsets within the CELLxGENE whole-human dataset. During the pretraining stage, an AdamW optimizer (Loshchilov and Hutter 2017) was utilized with a 0.0001 initial learning rate, which was decreased by a factor of 0.95 after each epoch. The model was trained for 8 epochs with a batch size of 32. For the tissue-specific client models, 90% of the data was used for training and the remaining 10% for validation. The corruption rate was set to 0.6 and the temperature coefficient of ℒ_contrast_ to 0.07. To achieve a relative balance between the two learning objectives, the weight of ℒ_contrast_ and ℒ_rec_ was set to 1:0.03. For the federated learning settings, local tabular transformer weight aggregation and update were performed at the end of every training epoch for all clients. Since all clients are considered equally important, the global tabular transformer weight was averaged across all clients during aggregation. After pretraining, we extract the client model weights following 8 epochs as the final tissue-specific embedders and project heads weights, and we use the aggregated weight of the transformer as the final global tabular transformer weight.
- Benchmark federated vs. centralized pretraining We benchmarked two pretraining schemes: federated learning and centralized learning. The federated learning scheme followed the same training methodology in Tabula. In contrast, the centralized learning scheme adopted a traditional approach, where all data was consolidated, and model pretraining was performed on a single centralized device using all acquired data. This approach avoids maintaining multiple models for individual clients and performing model weight aggregation and updates across clients.
- Benchmark tabular learning vs. MLM We benchmarked two single-cell modeling strategies: tabular learning and masked language modeling (MLM). The tabular learning strategy was using the same learning objectives in Tabula, while the MLM strategy aimed to reconstruct these masked expression values through contextual inference. Specifically, 15% of the non-zero gene expression values in the input sequence were randomly masked by setting their expression levels to -1, following scBERT (Yang et al. 2022). The mean squared error (MSE) loss is computed exclusively for the masked positions.
- Cell type annotation For the cell type annotation task, we adopted an initial learning rate of 0.0001 and a weight decay of 0.1. The batch size was set to 32, while finetuning was performed with an early stopping strategy. For the cell line dataset, we randomly splitted 33% of the data as the test set, with the remaining 67% used for training where 10% was randomly chosen as the validation set. In other datasets, the training and test sets were pre-defined (see **Downstream Task Datasets**), and 10% of the training set was similarly reserved as the validation set. The optimal weights were selected based on the lowest loss on the validation set and used for evaluating model’s performance on the test set. The data preprocessing steps included transcripts-per-million normalization, log1p transformation, and value binning. In addition, we selected 3,000 highly variable genes and prioritized the genes with non-zero expression in each cell during data loading until the maximum length requirement of 1,200 genes was reached. If fewer than 1,200 genes were selected, genes with zero expression will be filled. This approach ensures that the model senses more genes during the finetuning process while meeting the maximum length requirement.
- Multi-omics and multi-batch integration For multi-omics and multi-batch integration tasks, we selected 2,000 HVGs from the dataset and randomly chose 1,200 genes, which is the maximum input length of the pretrained Tabula, for each training epoch. The data preprocess steps included TPM normalization, log1p transformation, HVG selection and value binning. During finetuning, we used an initial learning rate of 0.0001 and a weight decay of 0.95. For datasets with multi-omics, the model was extended to accommodate new tokens for new omics modality similar to gene embeddings, the mask ratio of ℒ_MGM_ and ℒ_CMGM_ was set to 0.4, and the corruption rate of ℒ_reconstruction_ was set to 0.6. For datasets with multi-batch, ℒ_DA_ was adopted with the weight of 1. The four losses were applied to both tasks if the dataset has multiple batches and multiple omics. Gradient clipping with a value of 0.5 was enabled to avoid exploding gradients. The model was trained with an early stopping strategy. The best performance of each dataset is reported based on the best-combined validation loss.
- Genetic perturbation prediction For the gene perturbation prediction task, in order to reconstruct all genes to enable comprehensive understanding of inter-gene relationship, for the datasets exceeding the maximum input length of 1,200 genes, we randomly initialized the parts that could not be accommodated by the pretrained model weights. The model optimization adopted a learning rate of 0.0001 and a weight decay of 0.95. The batch size was set to 64, while the training was conducted with an early stopping strategy with patience of 5 epochs.
- Imputation For the imputation task, the initial learning rate was set to 0.0001 with a weight decay of 0.95. The batch size was set to 32, and fine-tuning was performed over a fixed 10 epochs. The maximum sequence length was set to 2,000. We masked the data to simulate dropout values. This process masked non-zero gene expression data for each cell, splitting it into training, validation, and test sets. For each cell, non-zero values were identified. If there were fewer than 5 non-zero values, no masking was applied. Otherwise, based on an exponential probability distribution (scale 20), 10% of the data was masked for validation and another 10% for testing. The masked values were set to 0, and the remaining data formed the training set. The data preprocessing steps included TPM normalization and log1p transformation. We only conducted a value binning preprocess for the gene expressions in the training and validation sets, and the test sets were used with log1p values. The model weights exhibiting the lowest loss on the validation set were selected as the optimal weights for evaluating performance on the test set.
- In silico gene regulatory network inference For the task of in silico gene regulatory network inference, tissue-specific embedders and project heads weights were applied across various biological systems. These included using blood-specific weight for hematopoiesis, pancreas-specific weight for pancreatic endogenesis, brain-specific weight for neurogenesis, and heart-specific weight for cardiogenesis. The data preprocess steps included TPM normalization, log1p transformation, HVG selection and value binning before the model inference. To capture the gene-gene relationships inherent in regulatory networks, an input length of 1,200 is insufficient. Consequently, we initialized Tabula with an extended input length and loaded pretrained weights that have compatible dimensions, specifically: 1,900 for pancreatic endogenesis dataset, 1,775 for hematopoiesis dataset, and 2,400 for both neurogenesis and cardiogenesis datasets.

## Downstream Task Datasets

- CELLxGENE The CELLxGENE dataset is a publicly accessible resource developed and maintained by the Chan Zuckerberg Initiative, designed for storing, exploring, and analyzing scRNA-seq data. It aggregates data from research projects across the globe. Through the CELLxGENE portal, we collect scRNA-seq data from approximately 40 million human cells, and the release version from July 25, 2023. After quality control, preprocessing and filtering, about 15 million human scRNA-seq data are obtained for Tabula pretraining. This comprehensive dataset includes a diverse range of 23,156 genes and 422 cell types from 133 tissues and 196 studies.
- PBMC 5K We use a high-throughput scRNA-seq dataset PBMC 5K from 10x Genomics containing samples of peripheral blood mononuclear cells (PBMCs) from healthy donors. These cells are tagged with a panel of TotalSeq-B antibodies (v3 chemistry). The dataset comprises a total of 5,247 cells and includes 33,538 genes.
- Jurkat The Jurkat cells are human T-lymphocyte cell lines derived from patients with acute T-cell leukemia and are widely used in immunology research. This dataset provides scRNA-seq data for Jurkat cells generated by the 10x Genomics. The dataset comprises a total of 3,258 cells and includes 32,738 genes.
- Melanoma The Melanoma dataset centres on scRNA-seq of WM989-A6-G3 melanoma cells using two technologies: DropSeq and Fluidigm’s C1 mRNA Seq HT chip(Torre et al. 2018). These methods allow a detailed analysis of the transcriptomic characteristics of melanoma cells. We select data sequenced using DropSeq technology. The dataset comprises a total of 8,640 cells and includes 32,287 genes.
- Adamson The Adamson perturbation is a scRNA-seq dataset derived from K562 human cell lines (Adamson et al. 2016), designed to study the effects of CRISPR interference on gene expression. The dataset contains RNA expression profiles for 65,337 raw cells, with 43,550 cells retained after quality filtering. It includes measurements for 32,738 genes across 87 genetic perturbations, following filtering from 90 initial perturbations. The dataset is preprocessed by GEARS data processing function (Roohani, Huang, and Leskovec 2023) for further finetuning.
- Norman The Norman perturbation dataset is also a scRNA-seq dataset generated from K562 human cell lines (Norman et al. 2019), designed to explore genetic interactions through CRISPR activation. The dataset captures RNA expression profiles from 111,668 raw cells, with 82,081 cells retained after quality filtering. It contains measurements for 33,694 genes across 105 single-gene perturbations and 131 dual-gene perturbations. The dataset is preprocessed by GEARS data processing function (Roohani, Huang, and Leskovec 2023) for further finetuning.
- Replogle The Replogle perturbation dataset uses a multiplexed CRISPR interference with single-cell transcriptome sequencing to analyze the effects of gene knockdowns across cell types, creating a comprehensive human cell genotype-phenotype atlas (Replogle et al. 2022). The dataset has been meticulously processed based on the curated protocols of scGPT (Cui et al. 2024), which includes a total of 171,542 samples, derived from 1,823 distinct one-gene perturbations, of which 99 target transcription factors. Additionally, the dataset features a test set consisting of 456 perturbations, with 25 specifically affecting transcription factors. The dataset is preprocessed by GEARS data processing function (Roohani, Huang, and Leskovec 2023) for further finetuning.
- Human Pancreas The Human pancreas dataset comprises data from five scRNA-seq studies on human pancreas cells, reprocessed by (J. Chen et al. 2023) for cell type annotation. In total, there are 14 cell types including alpha, beta, ductal, acinar, delta, pancreatic stellate, pancreatic polypeptide, endothelial, macrophage, mast, epsilon, Schwann, T cell and MHC class IIl. The reference set includes data from two sources containing 10,600 cells across 13 cell types, while the query data covers the remaining three sources consisting of 4,218 cells spanning 11 cell types. The query set includes MHC class II, absent in the reference set, but lacks macrophage, Schwann, pancreatic polypeptide and T cell, which are present in the reference set.
- Myeloid The Myeloid dataset provides a comprehensive analysis of subpopulation profiling of tumour-infiltrating myeloid cells at the pan-cancer level, by integrating single cell transcriptomic data from nine different cancer types (Baron et al. 2016). The dataset was randomly subsampled by (Cui et al. 2024). The reference set contains 9,748 cells from six cancer types including UCEC, PAAD, THCA, LYM, cDC2 and kidney, while the query set contains 3,430 cells from four cancer types including MYE, OV-FTC and ESCA.
- 293t & Jurkat cells The 293t & Jurkat cells dataset, generated using the 10x Genomics platform, profiles gene expression across 16,602 genes in two distinct cell lines: 293T cells, originating from human embryonic kidney and commonly used in gene expression and viral studies, and Jurkat cells, a T lymphocyte line derived from a leukemia patient and frequently employed in T cell signaling research. As reported by (Tran et al. 2020), the dataset comprises three experimental batches: Batch 1 consisting of 2,885 293T cells, Batch 2 containing 3,258 Jurkat cells, and Batch 3 featuring a balanced mixture of both cell types with 3,388 cells total distributed equally between 293T and Jurkat cell lines.
- Liver The Liver dataset contains a variety of human liver cell types from two primary sources. The reference dataset, processed by(MacParland et al. 2018), contains 8,444 parenchymal and non-parenchymal cells from five human livers. It includes 14 different cell types, including central venous sinusoidal endothelial cells, erythroid cells, hepatocytes, inflammatory and non-inflammatory macrophages, mature B cells, natural killer (NK) cells, periportal sinusoidal endothelial cells, plasma cells, portal endothelial cells, stellate cells, alpha-beta T cells, cholangiocytes and gamma-delta T cells. While the query dataset provided by(Aizarani et al. 2019). consists of 9,162 cells from normal human liver tissue, it overlaps with 7 of the cell types found in the reference dataset.
- 10x Multiome PBMC The 10x Multiome PBMC dataset captures dual-modality single cell profiles from human peripheral blood mononuclear cells through paired RNA and ATAC sequencing analysis. Processed by (Cao and Gao 2022), this dataset contains 9,631 cells derived from a healthy 25-year-old female donor, with granulocytes removed via cell sorting before nuclear isolation and sequencing. The data spans 29,095 genes for expression analysis and 107,194 regions for chromatin accessibility measurements. The cells are classified into 19 distinct immune populations, encompassing various T cell subtypes (CD4+ naive, CD4+ TCM, CD4+ TEM, CD8+ naive, CD8+ TEM 1, CD8+ TEM 2, MAIT, Treg, gdT), B cell subtypes (naive, intermediate, memory, plasma), myeloid cells (CD14+ monocytes, CD16+ monocytes, cDC, pDC), as well as NK cells and HSPCs.
- BMMC The BMMC dataset is a comprehensive multimodal single-cell profiling of bone marrow mononuclear cells from 12 healthy human donors (M. D. Luecken, Burkhardt, and Cannoodt 2021), originally designed for the Multimodal Single-Cell Data Integration Challenge at NeurIPS 2021. It includes paired single-cell RNA and protein abundance measurements across 12 batches. Processed by (Cui et al. 2024), the final dataset consists of 90,261 cells, with data on 13,953 genes and 134 surface proteins, along with detailed annotations for 45 immune cell subtypes.
- Perirhinal Cortex The Perirhinal Cortex dataset is a subset of a larger study by (Siletti et al. 2023), which created a comprehensive transcriptomic atlas of the human brain using over three million nuclei from around 100 dissections across various brain regions. From this dataset, two separate batches from the perirhinal cortex were selected for further analysis, following the methodology outlined by scGPT (Cui et al. 2024). The first batch contains 8,465 cells, while the second batch includes 9,070 cells. Together, the datasets comprise a total of 59,357 genes, with the ten unique cell types identified in the original study being used for analysis.
- DC This DC dataset contains scRNA-seq data from human blood dendritic cells (DCs), originally collected by (Villani et al. 2017) and processed by (Tran et al. 2020). It includes two batches, each with four distinct cell types: CD1C DC, CD141 DC, plasmacytoid DC (pDC), and double negative cells. To ensure non-identical cell compositions across batches, CD1C DCs are excluded from batch 1, and CD141 DCs are excluded from batch 2. Each batch contains 288 cells and 16,594 genes, with 96 pDCs and 96 double negative cells shared between the batches. Batch 1 includes 96 CD141 cells, while batch 2 includes 96 CD1C cells.
- Hematopoiesis The Hematopoiesis dataset is derived from (Qiu et al. 2022). The dataset consists of 1,947 cells representing 8 cell types across five developmental nodes. These cell types include: hematopoietic stem cells (HSC), megakaryocyte-erythrocyte progenitors (MEP), granulocyte-monocyte progenitors (GMP), monocytes (MON), neutrophils (Neu), megakaryocytes (Meg), erythrocytes (Ery), and basophils (Bas). Based on the known cell differentiation(Krumsiek et al. 2011), HSC first differentiates into MEP and GMP, with GMP further developing into Mon and Neu, while MEP diverts into Meg, Ery, and Bas.
- Cardiogenesis The Cardiogenesis dataset is obtained from the cardiogenic region of mouse embryos (de Soysa et al. 2019). The cardiogenic mesoderm differentiates from a shared population of cardiovascular progenitor cells into two regions with distinct gene expression: the first heart field (FHF) and the second heart field (SHF). We focus on the gene regulatory relationships within the FHF and SHF. A total of 1,165 cells are selected from this dataset for analysis, comprising cells from both the posterior second heart field (pSHF) and FHF.
- Neurogenesis The Neurogenesis dataset is a comprehensive single cell transcriptomic atlas of the embryonic mouse brain between gastrulation and birth (La Manno et al. 2021). We focus on the transformation of glial cells into astrocytic and oligodendrocytic lineages within the central nervous system. We select a total of 4,704 cells from the atlas, which include 2,516 mixed region astrocytes, 1,990 oligodendrocyte precursor cells, 162 committed oligodendrocyte precursor cells, and 36 mature oligodendrocytes.
- Pancreatic Endogenous The Pancreatic Endogenous dataset is based on the work of (J. Wang et al. 2020). We aim to investigate the master transcription factors that govern pancreatic cell fate during the development of the pancreas. A total of 14,445 cells are selected from the dataset across three time points: 4,631 cells at E12, 5,179 cells at E14, and 4,635 cells at E17.

**Supplementary Figure 1:**
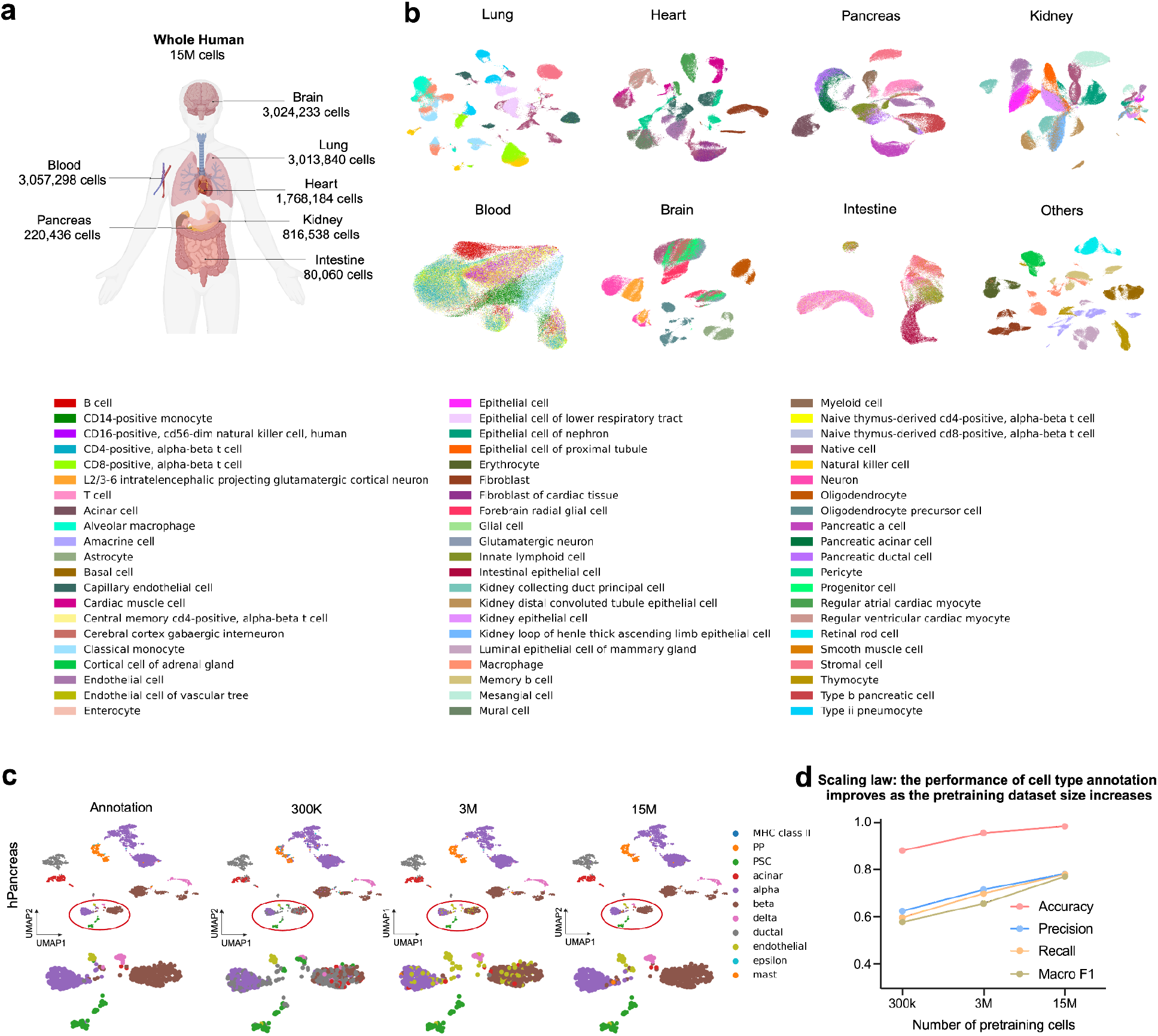
The overview of pretraining dataset and scaling law in Tabula. a. Summary of Tabula’s pretraining scRNA-seq data’s tissue type distribution in human. Tabula’s pretraining dataset includes around 15 million (M) scRNA-seq data distributed across many human tissues. To demonstrate the Tabula federated learning framework, we assign each tissue to a distinct client. b. UMAP visualization of cell embeddings from pretrained Tabula client models, with 10,000 cells randomly sampled from each client. The “Others” client aggregates all less-represented tissues. c. Tabula based pancreas-specific cell type annotation on the hPancreas dataset (J. Chen et al. 2023), visualized in the UMAP embedding. The performance improves as the size of the pretraining dataset increases from 300k to 3M, and to 15M cells. d. Scaling law: Impact of pretraining dataset size on the performance of cell type annotation in hPancreas dataset in panel **c** (evaluated by accuracy, precision, recall, and macro F1).

**Supplementary Figure 2:**
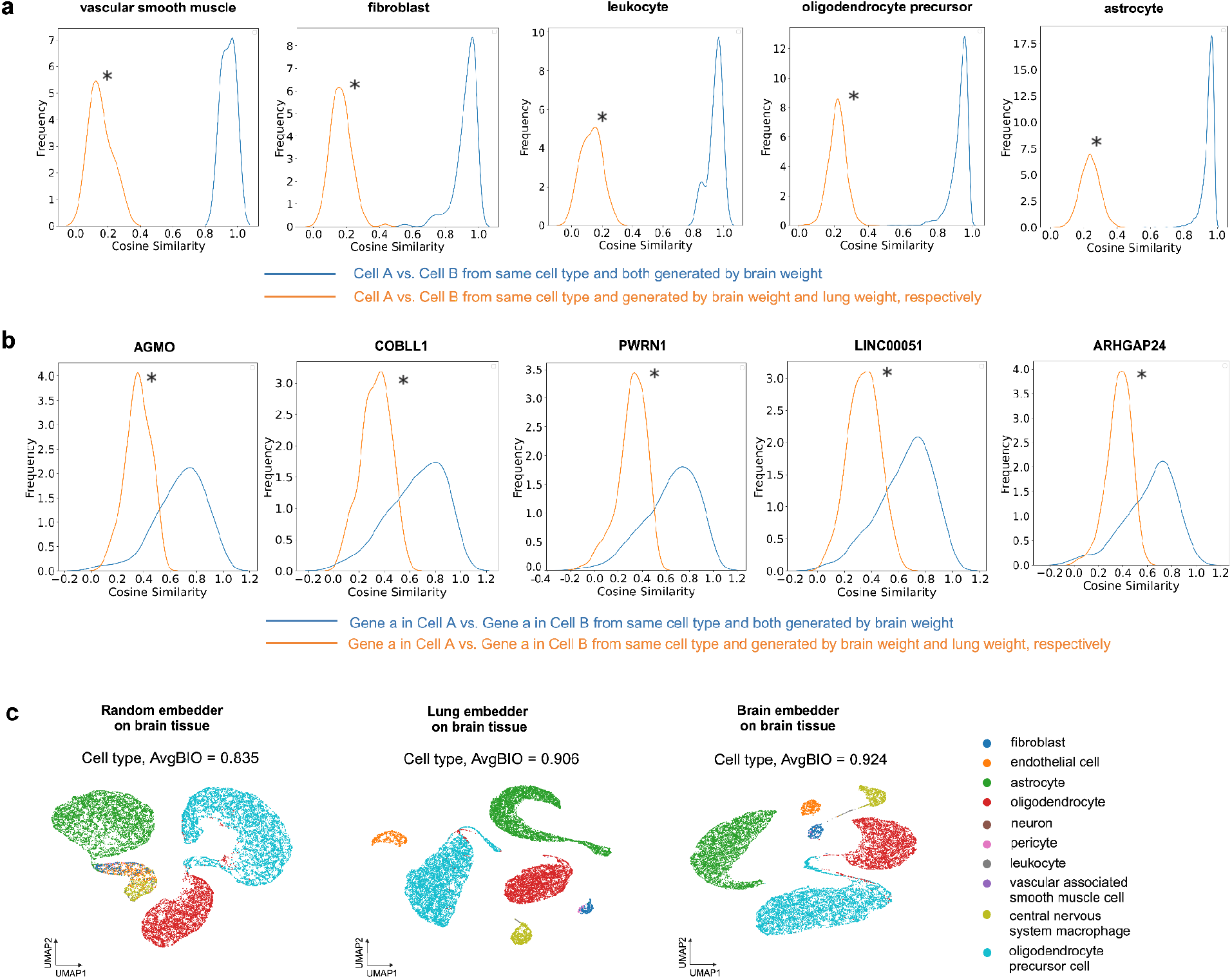
tissue-specific embedders encode distinctive tissue features. a. Tissue-specific embedding captures different biological contexts across various cell types at cell level. Perirhinal cortex dataset from brain tissue is adopted as an example. Each plot compares the cosine similarity between every possible cell pair from the same type both generated by brain-specific Tabula (line in blue) and between cells of the same type but derived from different tissues, brain and lung-specific Tabulaweights, respectively (line in orange). The distribution indicates the degree of similarity, with peaks close to 1 suggesting high intra-group consistency. The distribution from the blue lines is significantly shifted to the right (* indicates p < 0.05 by Wilcoxon test, FDR-corrected) highlights the tissue-specific Tabula models learn unique cellular molecular profiles of tissues. b. Same as in panel **a** but for comparing the cosine similarity of gene representations. c. Tissue-specific Tabula model demonstrates enhanced performance on corresponding datasets. By fine-tuning the Tabula model using diverse embedders on the Perirhinal cortex dataset derived from brain tissue, the brain-specific Tabula embedder notably outperforms embedders initialized with weights from other tissues and those with random initialization. This finding highlights the benefits of employing tissue-specific embedders in improving downstream tasks tailored to specific biological contexts.

**Supplementary Figure 3:**
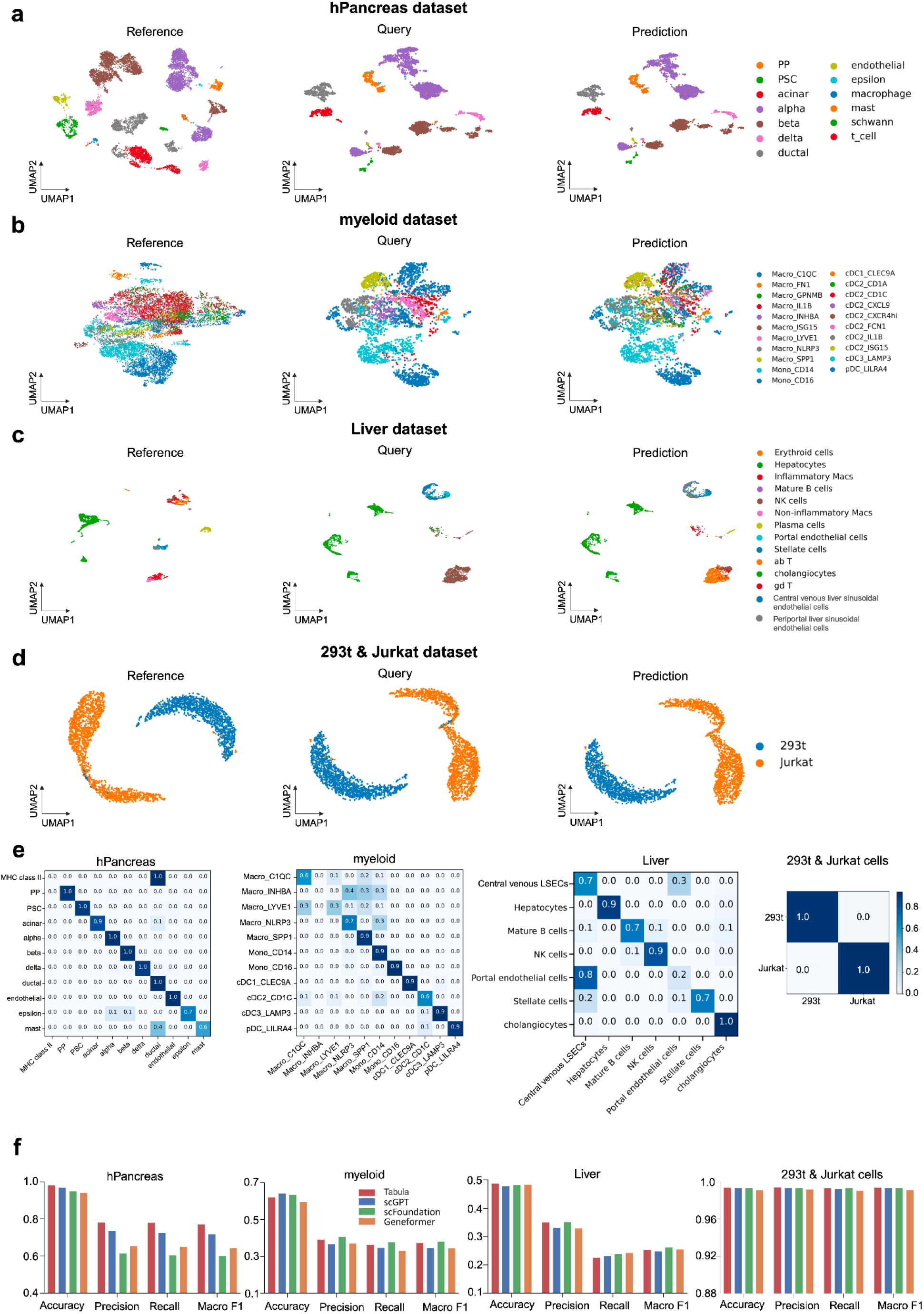
Tabula generally outperforms state-of-the-art single cell foundation model in annotating cell types. a. UMAP visualization of the hPancreas dataset, colored by cell types. The reference dataset (left) was used to fine-tune Tabula, which was evaluated on the query dataset (middle). The predicted annotations (right) demonstrate the model’s annotation capability. b. Similar to (a), but for the myeloid dataset. c. Similar to (a), but for the Liver dataset. d. Similar to (a), but for the 293t & Jurkat cells dataset. e. Confusion matrices for cell type annotation prediction of the hPancreas, Myeloid, Liver, and 293T & Jurkat cells datasets, revealing a majority of correct prediction along the diagonal. The x-axis represents the predictions, while the y-axis corresponds to the ground truth. f. Barplots of performance metrics (accuracy, precision, recall, Macro F1) of Tabula against scGPT, scFoundation, and Geneformer across the four datasets. Tabula consistently achieves superior or competitive results.

**Supplementary Figure 4:**
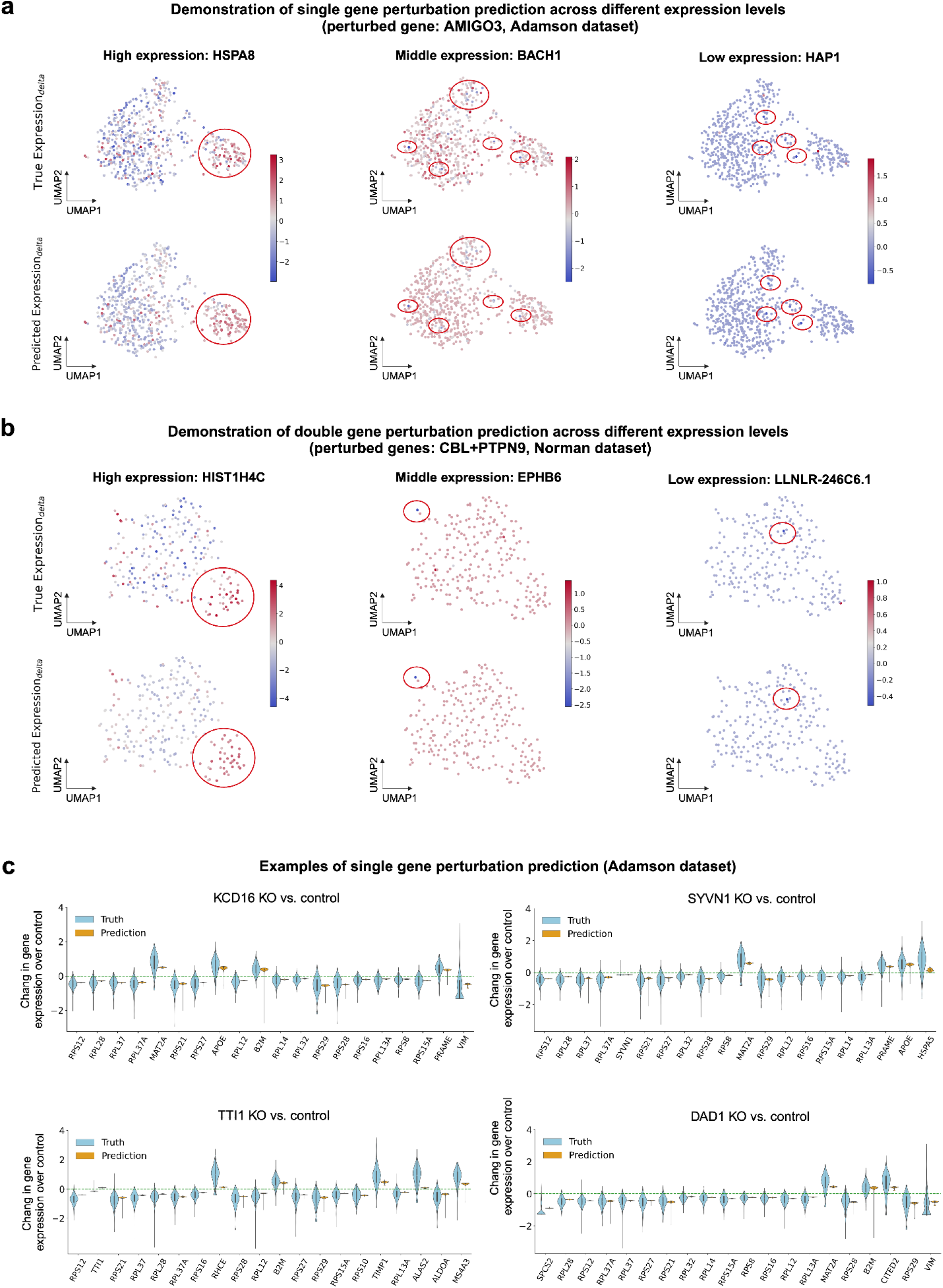
Scatterplot of target genes after single- or double-gene perturbation and examples of gene expression changes in response to single-gene perturbation. a. Scatterplots of example target genes with different expression levels after single-gene perturbation from the Adamson dataset (Adamson et al. 2016). The perturbed gene is AMIGO3, and the effects are displayed forthree target individual genes with different expression levels: high (HSPA8), medium (BACH1), and low (HAP1). b. Scatterplots of example target genes with different expression levels after double-gene perturbation from the Norman dataset (Norman et al. 2019). The perturbed genes are CBL and PTPN9, and the effects are displayed for three target individual genes with different expression levels: high (HIST1H4C), medium (EPHB6), and low (LLNLR-246C6.1). c. Comparison of the distribution of gene expression changes of top 20 DE genes from ground truth (blue) and Tabula’s predictions (orange) in response to perturbations of KCD16, SYVN1, TTI1, and DAD1 relative to control conditions (similar to **Fig. 4d**).

**Supplementary Figure 5:**
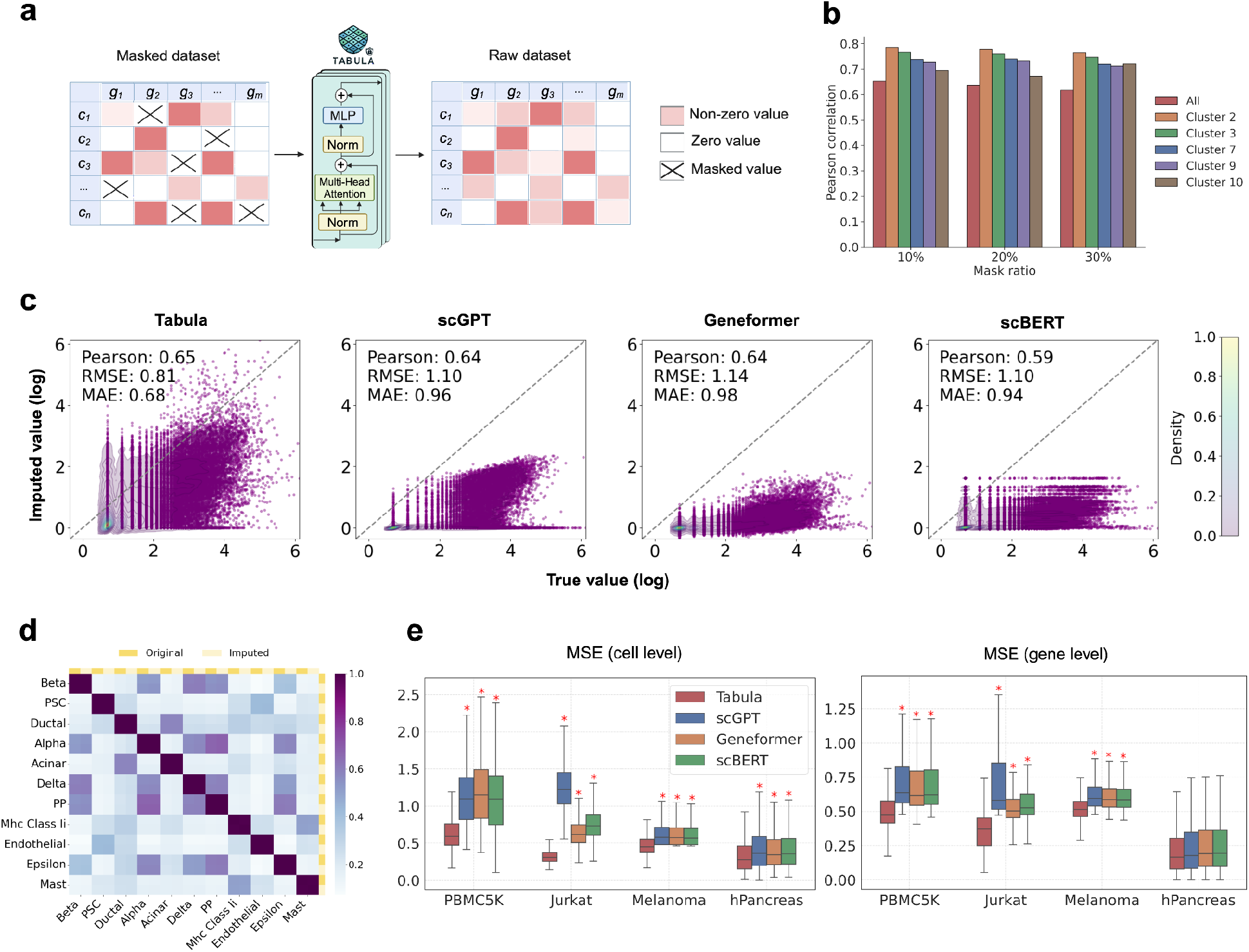
Tabula surpasses existing foundation models in gene expression imputation tasks across a variety of datasets. a. Schematic of the data imputation task: The dropout probabilities derived from an exponential distribution of gene expression are utilized to weight and sample non-zero values from a subset of genes within each cell, masking their expression to zero to simulate gene dropout. The fine-tuned Tabula model is subsequently applied to recover the true expression values of these masked genes. b. Robustness of the gene expression imputation by Tabula across varying gene masking ratios of 10%, 20%, and 30% on the PBMC5K dataset. The evaluation includes pearson correlation for both the overall dataset and the top five clusters. c. Scatterplot comparison of groundtruth (x-axis) vs. imputed values (y-axis) across Tabula, scGPT, Geneformer, and scBERT on the PBMC5K dataset. Log-transformed values are used for the density plot while the color of the points indicates the density of imputed values. Quantitative metrics, including Pearson correlation coefficient, Mean Absolute Error and Root Mean Square Error, are provided for each method. d. Fidelity and consistency of cell type similarity before and after Imputation. The heatmaps visualizes the degree of similarity between cell types in the original and imputed data on hPancreas dataset. Coloured squares at the edges represent the original and imputed data, while the varying colours within the heatmap illustrate the similarity between cell types. Tabula preserves the similarity between original and imputed data within the same cell type, while also maintaining the distinctions between different cell types. e. Boxplot of the mean squared error of the original and imputed values at both the cell-cell and gene-gene levels across four different datasets under four methods. Wilcoxon rank-sum test is used to determine the p-value between Tabula and other methods. P < 0.05 is denoted with an asterisk.

**Supplementary Figure 6:**
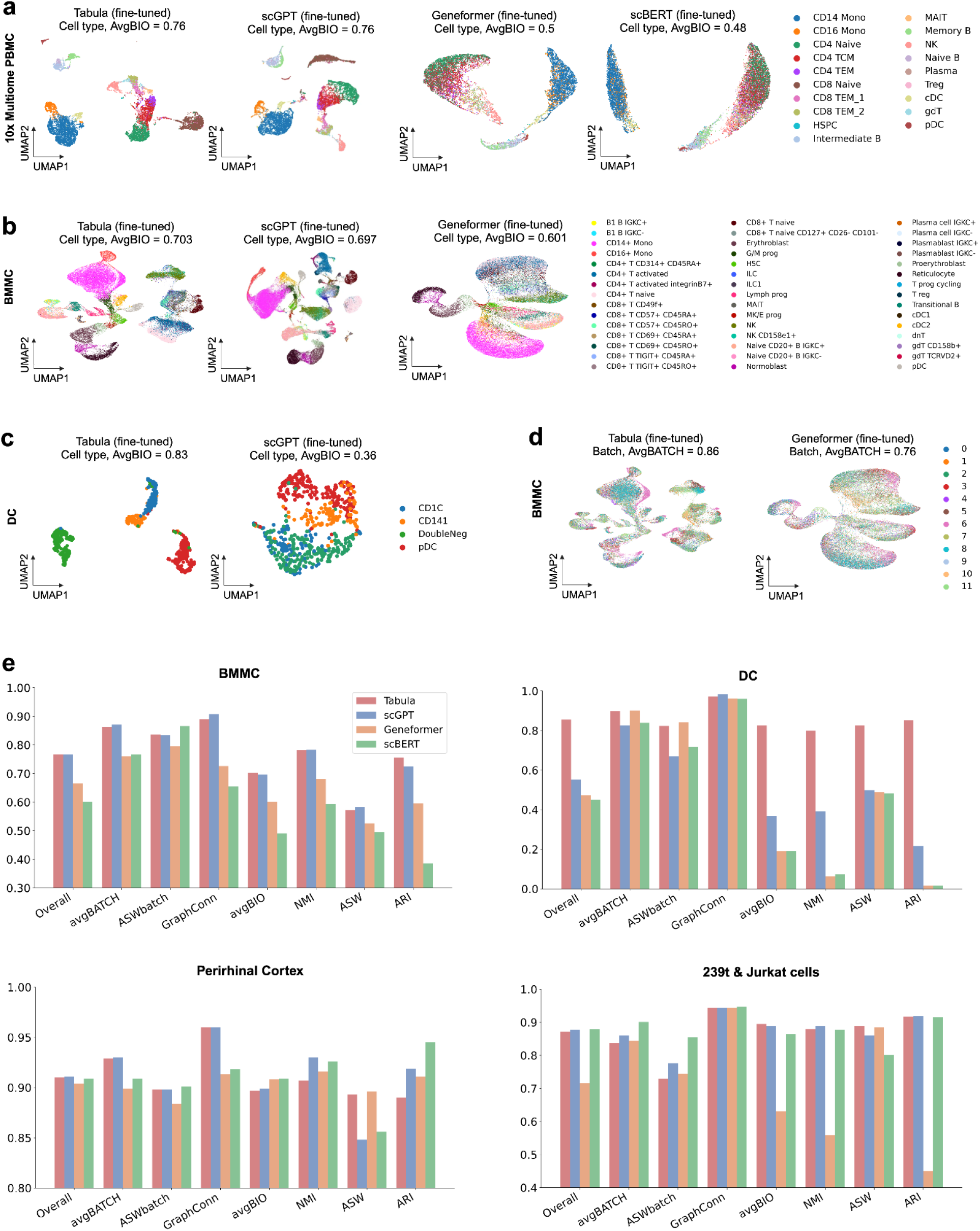
Multi-batch and multi-omics integration performance comparison. a. UMAP scatterplots of 10x PBMC Multiome dataset showing multi-omics integration by Tabula, scGPT, Geneformer, and scBERT. Tabula and scGPT exhibit superior biological integration performance (AvgBIO = 0.76) thanGeneformer and scBERT. b. Same as in panel **a** but for the BMMC dataset. Tabula demonstrates the highest AvgBIO (0.703), followed by scGPT (0.697) and then Geneformer (0.601). c. Same as in panel **a** but for the DC dataset. Tabula achieves a significantly higher AvgBIO score than scGPT. d. Same as in the panel **a** but for the BMMC dataset. Tabula achieves a significantly higher AvgBATCH score than Geneformer. e. Barplots of multi-omics and multi-batch integration metrics (AvgBATCH, ASWbatch, GraphConn, AvgBIO, NMI, ASW, ARI) across BMMC, DC, Perirhinal Cortex, and 293T & Jurkat cells datasets and under different models. Tabula consistently achieves top or competitive performance in biological and batch integration.

**Supplementary Figure 7:**
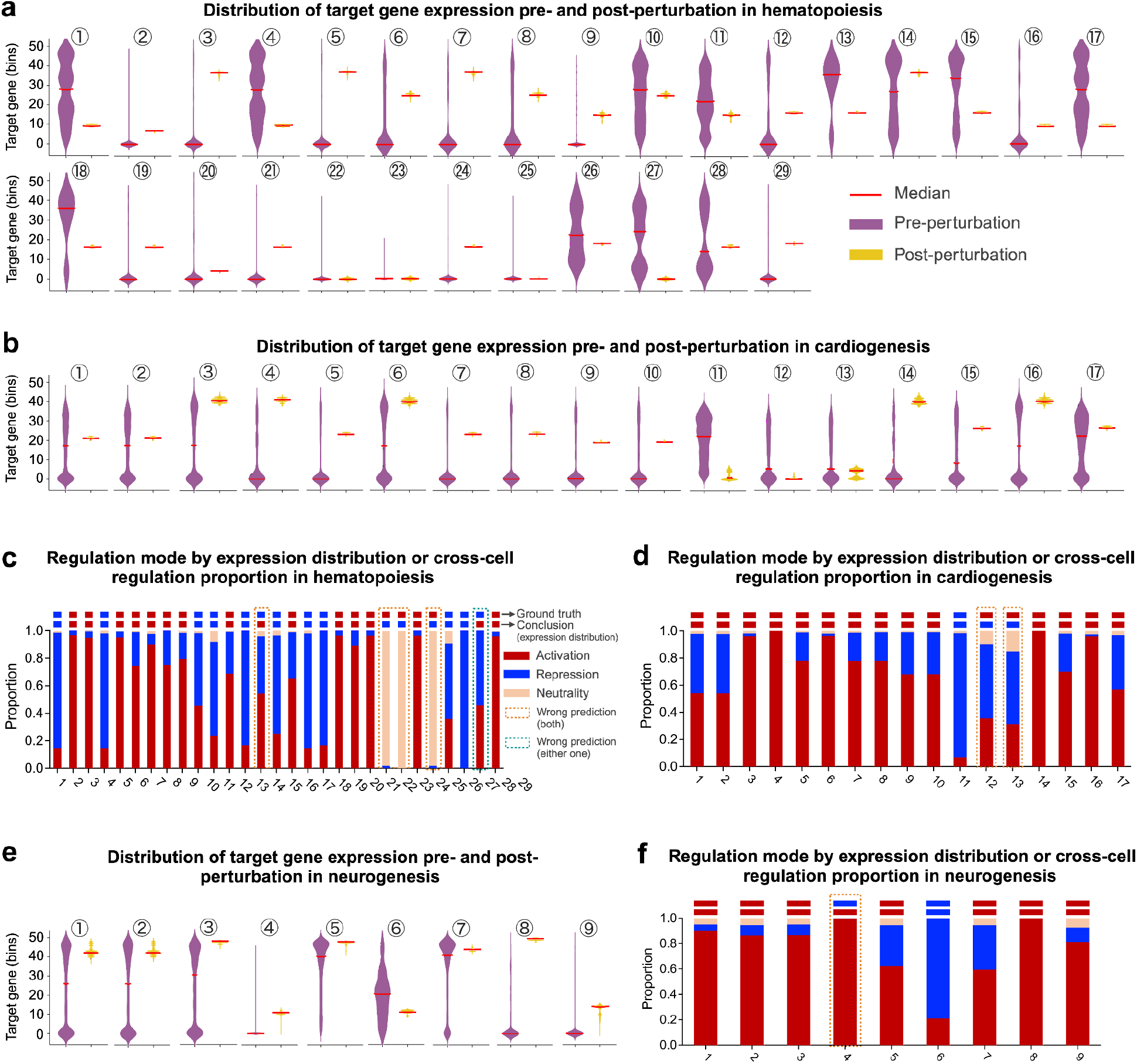
Pairwise regulation prediction by Tabula across hematopoiesis, cardiogenesis, and neurogenesis systems. a. Distribution of target gene expression pre- and post-perturbation under Tabula prediction within a network motif of 11 genes and 29 edges from the hematopoiesis system (Krumsiek et al. 2011) (similar to **Fig. 3d**). b. Same as in panel a but for the network motif of 9 genes and 17 edges cardiogenesis system (Nakanishi et al. 2016) (similar to **Fig. 3d**). c. Comparison of regulation mode identified by expression distribution shift and cross-cell regulation proportions in hematopoiesis system (Krumsiek et al. 2011) (similar to **Fig. 3e**). d. Same as in panel c but for the cardiogenesis system (Nakanishi et al. 2016). e. Same as in panel a but for the network motif with seven genes and nine edges from the neurogenesis system (Qiu, Ding, and Shi 2012). f. Same as in c but for the neurogenesis system (Qiu, Ding, and Shi 2012).

**Supplementary Figure 8:**
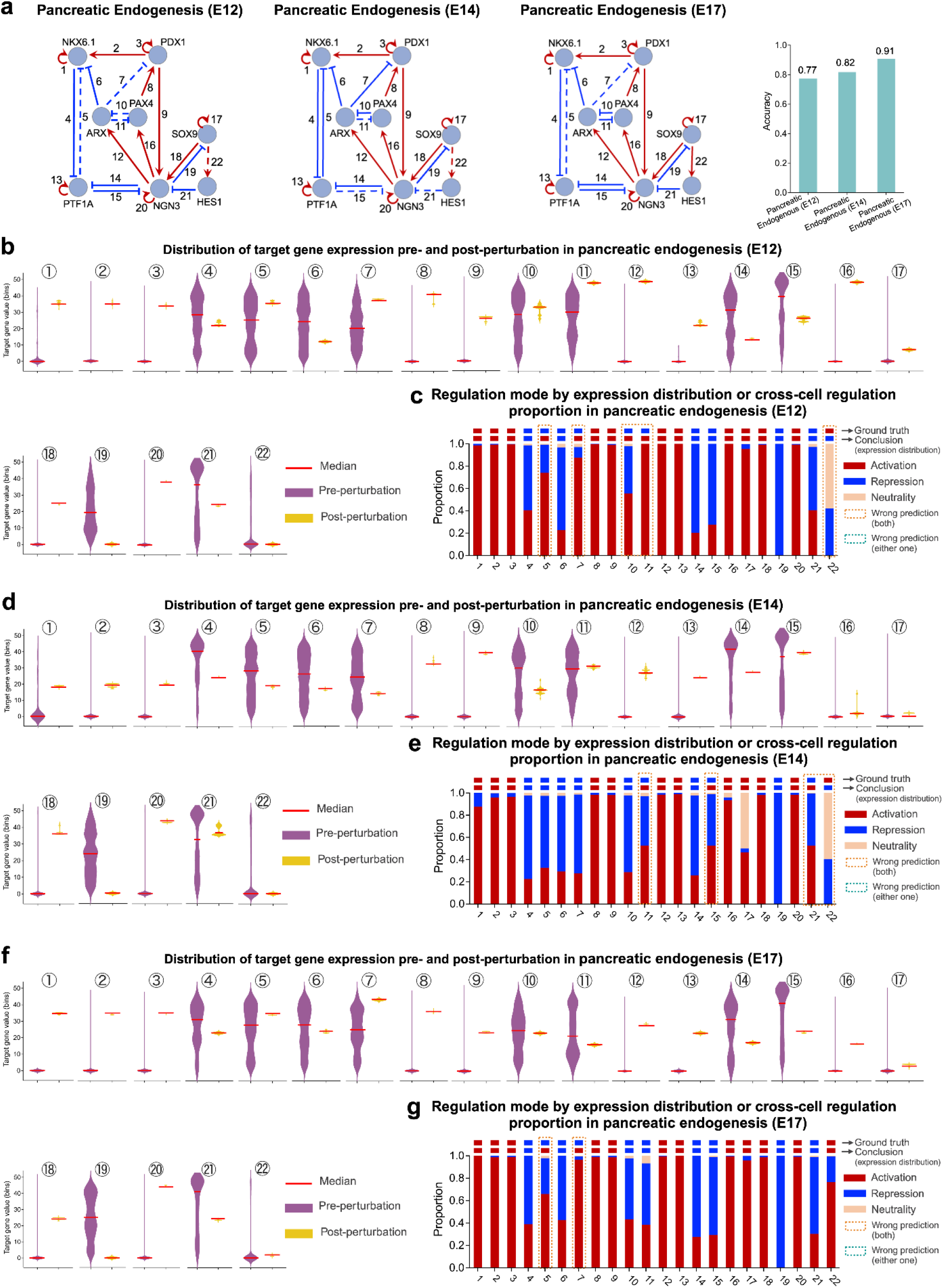
Pairwise regulation prediction by Tabula in pancreatic endogenesis system across developmental stages E12, E14, and E17. a. Pancreatic endogenesis system at three developmental stages (E12, E14, E17) with extensively validated regulatory networks that Tabula applied to and its performance in pairwise regulation prediction (J. Wang et al. 2020) (similar to **Fig. 3b**). b. Distribution of target gene expression pre- and post-perturbation under Tabula prediction in the pancreatic endogenesis core network at developmental stage E12 featuring 22 edges (J. Wang et al. 2020) (similar to **Fig. 3d**). c. Comparison of regulation mode identified by expression distribution shift and cross-cell regulation proportions in pancreatic endogenesis system at developmental stage E12 (J. Wang et al. 2020) (similar to **Fig. 3e**). d. Same as in panel b but for the developmental stage E14. e. Same as in panel c but for the developmental stage E14. f. Same as in panel b but for the developmental stage E17. g. Same as in panel c but for the developmental stage E17.

**Supplementary Figure 9:**
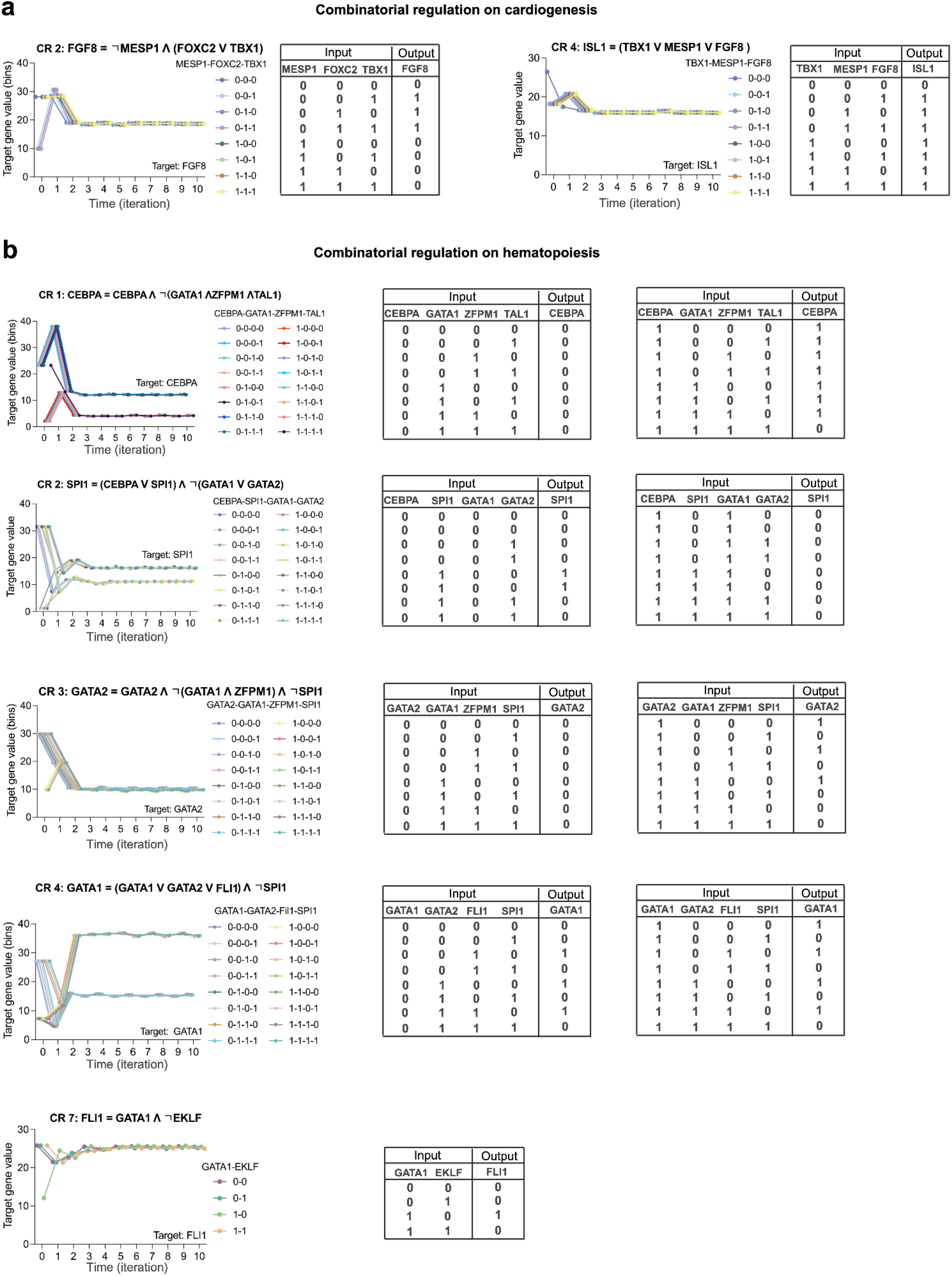
Combinatorial regulation predictions by Tabula for cardiogenesis and hematopoiesis systems. a. Combinatorial regulation (CR) predictions of CR2 and CR4 for the cardiogenesis system (Herrmann et al. 2012) (similar to **Fig. 3f**). b. Combinatorial regulation (CR) predictions of CR1-4 and CR7 for the hematopoiesis system (Krumsiek et al. 2011) (similar to **Fig. 3g**).

## SUPPLEMENTARY NOTES

### S1 Evaluation metrics for cell type annotation

To evaluate the performance of cell type annotation tasks, we use standard classification metrics: accuracy, precision, recall, and the F1 score. These metrics are defined as follows:

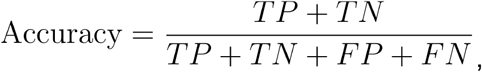

where *TP* is the number of true positives, *TN* is the number of true negatives, *FP* is the number of false positives, *FN* and is the number of false negatives.

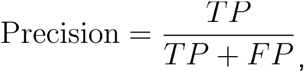

where precision quantifies the proportion of correctly predicted positive cell types among all predicted positives.

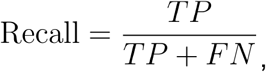

where recall measures the ability to identify all true positive instances of a cell type.

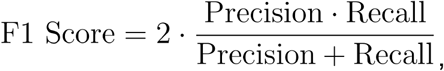

where F1 score provides a balanced measure that accounts for both precision and recall, especially useful for imbalanced datasets.

### S2 Evaluation metrics for multi-omics and multi-batch integration

To evaluate the performance of integration tasks, we adopt the widely used evaluation pipeline in the field (Malte D. Luecken et al. 2022) and selectively choose five metrics as conducted by scGPT (Cui et al. 2024). For multi-omics integration, metrics that measure conservation of biological variance of cell type, including, *NMI*_*cell*_, *ARI*_*cell*_ and *ASW*_*cell*_, are considered. For multi-batch integration, graph connectivity (*Graph*_*conn*) and *ASW*_*batch*_ are used for evaluation.

NMI (Normalized Mutual Information) measures the agreement between predicted and true cluster labels, providing an indication of how well cell types are preserved post-integration. It ranges from 0 (no agreement) to 1 (perfect agreement).

ARI (Adjusted Rand Index) evaluates the similarity between clustering results and true cell type annotations, adjusted for random chance. A higher ARI indicates better preservation of biological groupings.

ASW (Average Silhouette Width) quantifies the separation of cells within their assigned clusters compared to other clusters. A higher ASW indicates better-defined and more biologically meaningful clusters.

The batch-specific ASW evaluates how well batch effects are removed while maintaining meaningful biological groupings. A lower batch ASW indicates better integration across different batches.

Graph connectivity measures the continuity of the integrated data in a shared embedding space. Higher connectivity reflects improved integration, ensuring that cells from the same biological group form a cohesive cluster regardless of batch origin.

For systematically assessing the performance of integration tasks, *AvgBIO* and *Avg*_*batch*_ are employed to evaluate multi-omics integration and multi-batch integration, respectively. Additionally, *Overall* is used to measure the comprehensive performance of both tasks when the dataset is used to simultaneously evaluate the two tasks. The formulas are defined as follows:

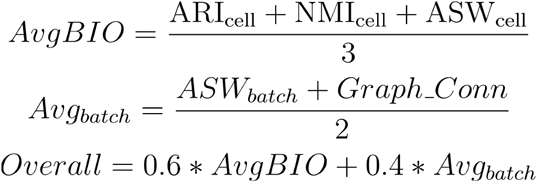

### S3 Evaluation metrics for genetic perturbation prediction

To evaluate the performance of genetic perturbation prediction tasks, we use Pearson correlation coefficient, which quantifies the linear relationship between predicted and actual expression changes following perturbation. The Pearson coefficient computed across all predicted gene expression changes for a cell is formulated as follows:

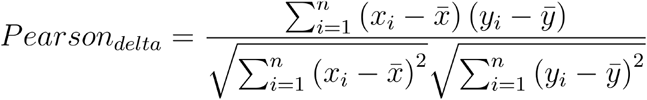

Here, *x*_*i*_ and *y*_*i*_ represent the predicted and actual gene expression changes for gene *i*, respectively. 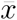 and 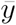 are the mean values of the predicted and actual expression changes across all genes. is the *n* total number of genes.

Similar to *Pearson*_*delta*_, the Pearson coefficient computed across top 20 differentially expressed gene expression changes for a cell, *DEG Pearson*_*delta*_, is formulated as follows:

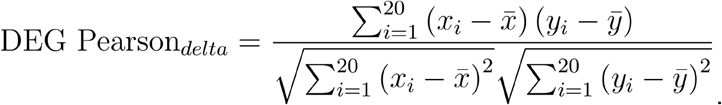

Here, *x*_*i*_ and *y*_*i*_ represent the predicted and actual gene expression changes for the top 20 differentially expressed genes (DEGs), respectively.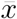 and 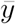 denote the mean values of the predicted and actual expression changes for these DEGs.

### S4 Evaluation metrics for imputation

To evaluate the performance of imputation tasks, we use multiple metrics, including the Pearson correlation coefficient, Root Mean Squared Error (RMSE), and Mean Absolute Error (MAE). The Pearson correlation coefficient quantifies the linear relationship between imputed and true gene expression values, while RMSE and MAE measure the differences between them. all metrics are computed across all imputed gene expression values and formulated as follows:

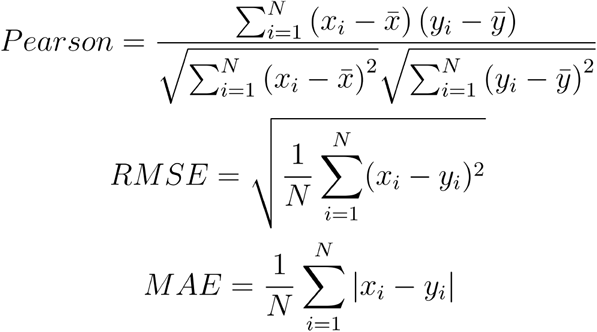

Here, *x*_*i*_ and *y*_*i*_ represent the imputed and true gene expression values for masked gene, respectively. 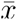 and 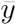 are the mean values of the imputed and true gene expression values across all masked genes. *N* denotes the total number of masked genes across all cells.

To evaluate the consistency between the original and imputed data, we calculate the Mean Squared Error (MSE) at two levels: cell-cell and gene-gene. The definitions are as follows: *MSE*_*cell*-*cell*_ measures the differences between the imputed and true values for all masked genes within a given cell. The formula is:

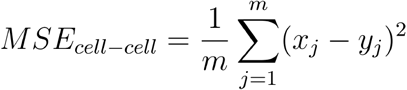

Here, *x*_*j*_ and *y*_*j*_ represent the imputed and true gene expression values for masked gene in a given cell. *m* is the total number of masked genes in this cell.

Similar to *MSE*_*cell*-*cell*_, *MSE*_*gene*-*gene*_ measures the differences between the imputed and true gene expression values for all masked cells within a given gene. The formula is:

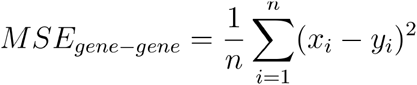

Here, *x*_*i*_ and *y*_*i*_ represent the imputed and true gene expression values for masked cell in a given gene. *n* is the total number of masked cells in this gene.

